# Dual Dysfunction of Kir2.1 Underlies Conduction and Excitation-Contraction Coupling Defects Promoting Arrhythmias in a Mouse Model of Andersen-Tawil Syndrome Type 1

**DOI:** 10.1101/2021.06.17.448833

**Authors:** Álvaro Macías, Andrés González-Guerra, Ana I. Moreno-Manuel, Francisco M. Cruz, Nieves García-Quintáns, Lilian K. Gutiérrez, Marta Roche-Molina, Francisco Bermúdez-Jiménez, Vicente Andrés, María Linarejos Vera-Pedrosa, Isabel Martínez-Carrascoso, Juan A. Bernal, José Jalife

## Abstract

Andersen-Tawil Syndrome (ATS) is associated with life threatening arrhythmias of unknown mechanism. We report on a mouse model carrying the trafficking-deficient mutant Kir2.1^Δ314-315^. The mouse recapitulates the electrophysiological phenotype of type 1 (ATS1), with slower conduction velocities in response to flecainide, QT prolongation exacerbated by isoproterenol, and increased vulnerability to calcium-mediated arrhythmias resembling catecholaminergic polymorphic ventricular tachycardia (CPVT). Kir2.1^Δ314-315^ expression significantly reduced inward rectifier K^+^ and Na^+^ inward currents, depolarized resting membrane potential and prolonged action potential duration. Immunolocalization in wildtype cardiomyocytes and skeletal muscle cells revealed a novel sarcoplasmic reticulum (SR) microdomain of functional Kir2.1 channels contributing to intracellular Ca^2+^ homeostasis. Kir2.1^Δ314-315^ cardiomyocytes showed defects in SR Kir2.1 localization and function, which contributed to abnormal spontaneous Ca^2+^ release events. This is the first in-vivo demonstration of a dual arrhythmogenic mechanism of ATS1 defects in Kir2.1 channel function at the sarcolemma and the SR, with overlap between ATS1 and CPVT.

## INTRODUCTION

Trafficking deficient mutations in the gene coding the strong inward rectifier K^+^ channel Kir2.1 (*KCNJ2*) result in the autosomal-dominant Andersen-Tawil syndrome type 1 (ATS1)^1, 2^. ATS1 is manifested as a triad of ventricular arrhythmias, periodic paralysis and dysmorphic features^3, 4^. Its electrocardiographic manifestations include QT prolongation, ventricular ectopy, bigeminy, polymorphic or bidirectional ventricular tachycardia and sudden cardiac death (SCD) ^5^. In some “atypical cases” ATS1 overlaps phenotypically with catecholaminergic polymorphic ventricular tachycardia (CPVT) ^5, 6^. Kir2.1 is the main channel controlling both the resting membrane potential and the final phase of ventricular repolarization ^7^. Together with the main cardiac voltage-gated Na^+^ channel (NaV1.5), Kir2.1 forms channelosomes that help to control ventricular excitability and action potential propagation velocity ^7, 8^. Emerging evidence indicates that these two widely different channels physically interact from early stages of protein assembly and trafficking, and they share common partners that may include, but are not limited to, anchoring/adapter proteins, enzymes, and scaffolding and regulatory proteins ^8–12^, highlighting the relevance of macromolecular ion channel interplay in cardiac diseases ^13^.

*In vitro* approaches have demonstrated that NaV1.5 and Kir2.1 proteins colocalize at the cell membrane and regulate each other’s levels via PDZ-mediated binding to either the MAGUK protein SAP97 or α1-syntrophin, both of which help to stabilize both channels at the cell membrane ^8, 9^. Moreover, co-expression of trafficking deficient mutant Kir2.1^Δ314-315^ channels with wildtype (WT) NaV1.5 reduces both *I_K1_* and *I_Na_*, suggesting that the NaV1.5-Kir2.1 complex preassembles early in the forward trafficking pathway, and that both channels traffic more efficiently as the NaV1.5-Kir2.1 complex than alone ^9, 10, 14^. However, it is unknown whether trafficking-deficient mutations *in vivo* affect NaV1.5-Kir2.1 interactions leading to reduced excitability and ventricular arrhythmias severe enough to result in SCD. Also unknown is the mechanism by which some *KCNJ2* mutation carriers present arrhythmias resembling CPVT.

We postulate that in addition to reduced *I_K1_*, a proportion of ATS1 patients have reduced *I_Na_* in the atria and ventricles, because of trafficking disruption of the macromolecular complex that includes Kir2.1 and NaV1.5, leading to conduction defects and arrhythmias. In addition, we present new evidence indicating that, in addition to Kir2.1 channels that co-localize with NaV1.5 at the sarcolemma to control excitability, a unique population of sarcoplasmic reticulum (SR) Kir2.1 channels may help to maintain intracellular Ca^2+^ homeostasis and excitation-contraction (e-c) coupling. If oriented as we surmise, SR Kir2.1 channels should underlie a fundamental countercurrent for SR Ca^2+-^ATPase (SERCA)-mediated Ca^2+^ reuptake to the ryanodine receptor (RyR) mediated Ca^2+^ release. In addition, we demonstrate herein that defects in SR Kir2.1 localization and function contribute to abnormal spontaneous Ca^2+^ release in ATS1. Altogether, our analysis reveals a novel dual molecular mechanism at the sarcolemma and the SR membrane for the characteristic life-threatening arrhythmias in ATS1 patients, and how SR Kir2.1 channel dysfunction contributes to the phenotypic overlap between ATS1 and CPVT.

## RESULTS

### Kir2.1^Δ314-315^ is a trafficking deficient Kir2.1 mutant protein *in vivo*

Using AAV-mediated gene transfer, we have generated homogeneous cohorts of mice expressing the WT *KCNJ2* gene coding the inward rectifier K^+^ channel Kir2.1, and a *KCNJ2* gene variant coding a trafficking deficient protein with an internal deletion (Δ314-315)^10, 15^ called Kir2.1^Δ314-315^ (**Figure 1A**). We achieved cardiomyocyte-specific gene expression in live mice by i.v. AAV serotype 9 gene delivery in combination with shuttle constructs expressing target genes under the transcriptional control of the troponin-T proximal promoter (*cTnT)* as described ^16^. Test mice transduced with a reporter construct encoding Luciferase were used to demonstrate specific and homogeneous distribution throughout the ventricles (**Figure 1B**). Using the same cardio-specific shuttle AAV vector we expressed *in vivo* wild-type Kir2.1 (*Kir2.1WT*) or *Kir2.1^Δ314-315^* mutant followed by enhanced tandem dimer tomato (tdTomato) fluorescence protein after an internal ribosome entry site (IRES). A single i.v. injection of AAV resulted in a long-lasting, and homogeneous gene transexpression in cardiomyocytes (**Figure 1C, D**), without detectable expression in other cellular heart components like fibroblasts (**Online Figure I**). Efficiency of cardiomyocyte transduction after viral infection was measured by tdTomato immunofluorescence. Quantitative analysis revealed that more than 90% of the cells carried between 1 and 3 viral genomes (**Figure 1E**). Thus, AAV Kir2.1 transduction resulted in a 50% increase on total Kir2.1 protein level (**Figure 1F**), very much on the physiological range. As expected from a trafficking deficient mutant which is retained at the Golgi apparatus^10^, biochemical analysis showed less Kir2.1^Δ314-315^ protein transported to the plasma membrane compared to mice transduced with the WT isoform (**Figure 1G**) confirming a trafficking defect *in vivo* in a cardiac context.

**Figure 1.**
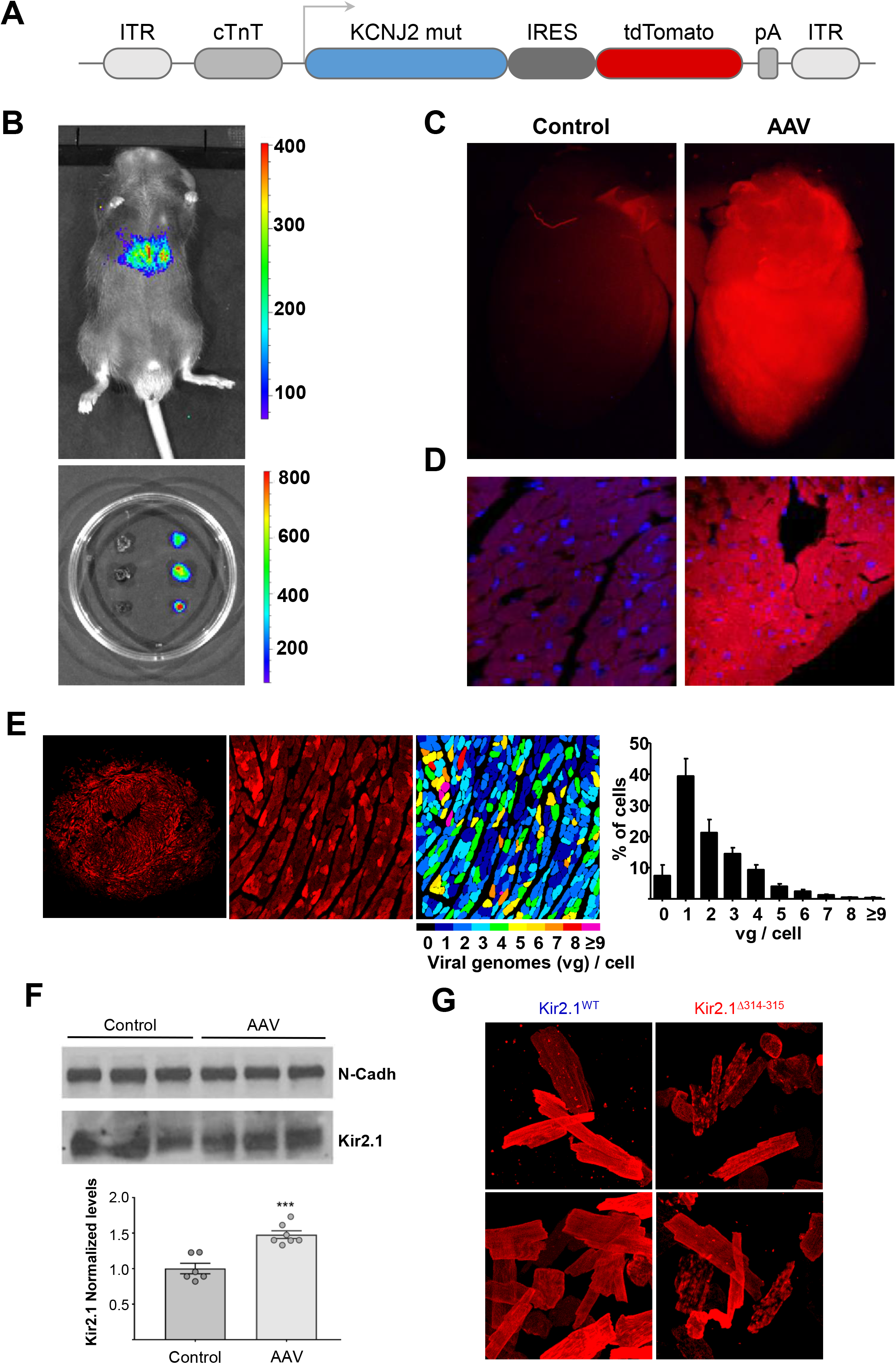
AAV-mediated *in-vivo* expression of Kir2.1^Δ314-315^. **A.** AAV based vector encoding human *KCNJ2* 314-315 mutant driven by the cardiac troponin-T (*cTnT*) proximal promoter, followed by enhanced tdTomato after an internal ribosome entry site (IRES). **B**, Imaging of luciferase transgene expression under *cTnT* cardiospecific control. *Upper*, live C57BL6J mouse injected (femoral vein) with AAV–based Kir2.1^Δ314-315^ vector in packaging serotype 9 (dose 3.5×10^10^ viral genomes [vg] in 50 μl of saline solution). *Lower*, three isolated hearts from Kir2.1^Δ314-315^ injected animals are shown next to 3 hearts from mock-injected mice (saline solution). Images taken 4 weeks after inoculation. **C-D.** Representative fluorescence images of 2 mock-injected (left) and 2 AAV-transduced mice (right), the latter showing expression of tdTomato throughout the heart. **E.** *Left,* representative fluorescence microscopy images of short-axis cross sections of AAV-transduced hearts illustrate expression of tdTomato throughout the heart. *Middle*, a cropped image used for quantification. *Right,* fluorescence intensity staining and quantification (N=3 mice, n=1465 cells) of transduced protein expression, used to assign the number of integrated viral genomes per cardiomyocyte. **F.** Representative western blots and quantification of total and membrane protein extracts from control and AAV-transduced isolated cardiomyocytes. Levels of expression were corrected by N-Cadherin (N-Cadh) expression and normalized by mock-injected levels. **G.** Representative immunostaining pattern of Kir2.1 at the plasma membrane in isolated cardiomyocytes from Kir2.1^WT^ and Kir2.1^Δ314-315^ expressing mice. Each value is represented as mean ± SEM. Statistical analyses were conducted using two-tailed *t*-test. ***, p<0.001.

### The Kir2.1^Δ314-315^ mouse model recapitulates the electrophysiological phenotype of ATS1 patients

Analysis of the resting ECG reveales that Kir2.1^Δ314-315^ mice suffer sinus arrhythmia and frequent premature ventricular contractions (PVCs) in the form of bigeminy (**Figure 2A**). These electrical abnormalities produced by mutant transgene expression were independent of any structural or functional defect as assessed by echocardiography and CMR imaging in AAV-injected animals (**Online Figure II**). ECG analysis also revealed significantly longer QTc intervals in Kir2.1^Δ314-315^ mice compared with Kir2.1^WT^ controls, while the QRS and PR intervals were similar in both groups (**Figures 2A-D;** time 0). To unmask and further characterize the electrical phenotype of ATS1 mice, we conducted a flecainide challenge experiment. Administration of a single i.p. dose of the drug led to progressively prolongation in both QRS and PR intervals (**Figure 2B**). After flecainide administration, the QRS duration in the Kir2.1^Δ314-315^ mice was more than twice that in the Kir2.1^WT^ mice, as would be expected by impaired trafficking of the Kir2.1-NaV1.5 channelosome produced by the 314-315 internal deletion on Kir2.1 protein ^10^. Furthermore, we investigated the incidence of arrhythmias induced by isoproterenol (ISO) administration. Kir2.1^Δ314-315^ mice showed increased arrhythmogenesis in response to ISO with obvious repolarization abnormalities, occasional U waves, ventricular extrasystoles (black arrows in **Figure 2C**), and a rapid increase in the QTc interval, without significant changes in the PR interval (**Figure 2D**). Kir2.1^Δ314-315^ animals were also more vulnerable than control animals to AF and VT/VF induced by endocardial burst pacing or the S1-S2 protocol (**Figure 2E**). Arrhythmia vulnerability was already present in resting conditions (**i**), but was especially evident after flecainide (**ii**) or ISO (**iii**) administration. In some cases, VT was polymorphic and converted to VF (**Figure 2E)**. Altogether, the foregoing data demonstrate that arrhythmias in the AAV-mediated ATS1 mouse model are a consequence of reduced cardiac excitability and conduction which are aggravated by stress.

**Figure 2.**
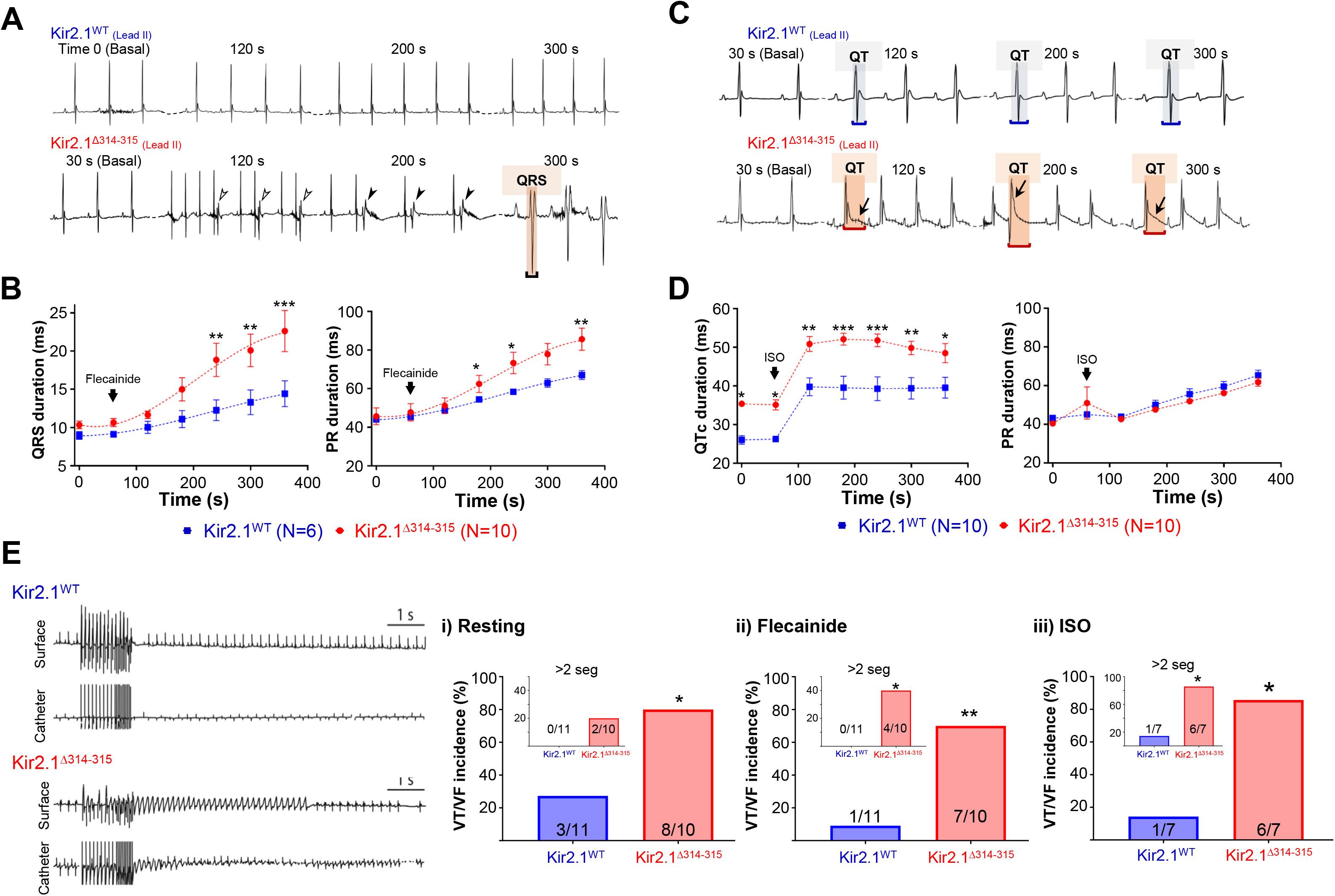
The Kir2.1^Δ314-315^ mouse model reproduces the electrical phenotype of Andersen-Tawil syndrome. **A** and **C.** Representative ECG recordings in Kir2.1^WT^ (top) and Kir2.1^Δ314-315^ (bottom) anesthetized mice showing the different ECG pattern in each group at different time points after a single dose of flecainide (20 mg/kg) and isoproterenol (isoprenaline, ISO, 100 mg/Kg), respectively. **B.** Quantification of temporal changes in QRS and PR interval durations before and after flecainide. Every value represents ten averaged intervals of ten consecutive beats. **D.** Corrected QT and PR intervals before and after ISO dose. Each value represents ten averaged intervals from ten consecutive beats. Data are expressed as mean ± sem. **E**. Representative Lead-II ECG traces and corresponding intracardiac recordings from anesthetized Kir2.1^WT^ and Kir2.1^Δ314-315^ mice. Graphs show the incidence of ventricular tachycardia/fibrillation (VT/VF) in resting conditions (i), after flecainide (ii) and after ISO (iii). Animals that had at least one AF episode ≥2 sec after burst pacing or S1-S2 stimulation were represented as graph insets. A period of polymorphic ventricular tachycardia (PVT) is shown. Each value is represented as mean ± SEM. Statistical analyses were conducted using two-tailed t-test and Chi-square Fisher’s exact test. *, p<0.05; **, p<0.01; ***, p<0.001

### Kir2.1-NaV1.5 channelosome function is impaired in ATS1 mice

Patch-clamp experiments demonstrated that, in accordance with previous *in-vitro* reports^10^, cardiomyocytes from Kir2.1^Δ314–315^ mice had reduced IK1 compared to cardiomyocytes from WT mice (**Figure 3A**). We also determined whether the expression of the trafficking-deficient Kir2.1^Δ314–315^ channel modified the cardiac INa. As illustrated in **Figure 3B**, cardiomyocytes from AAV-Kir2.1Δ314–315 mice exhibited a ~35% reduction in sodium current density compared to cardiomyocytes from mice transduced with Kir2.1^WT^. However, such differences were not due to differences in their voltage-dependence of activation or inactivation (**Online Figure III**). Our data would predict that in current-clamp assays the resting membrane potential of Kir2.1^Δ314–315^ would be more depolarized than control. Indeed, this is the case as only in 1 out of 9 cells (~11%) were we able to generate evoked action potentials, but with substantially longer duration than WT (**Figure 3C**).

**Figure 3.**
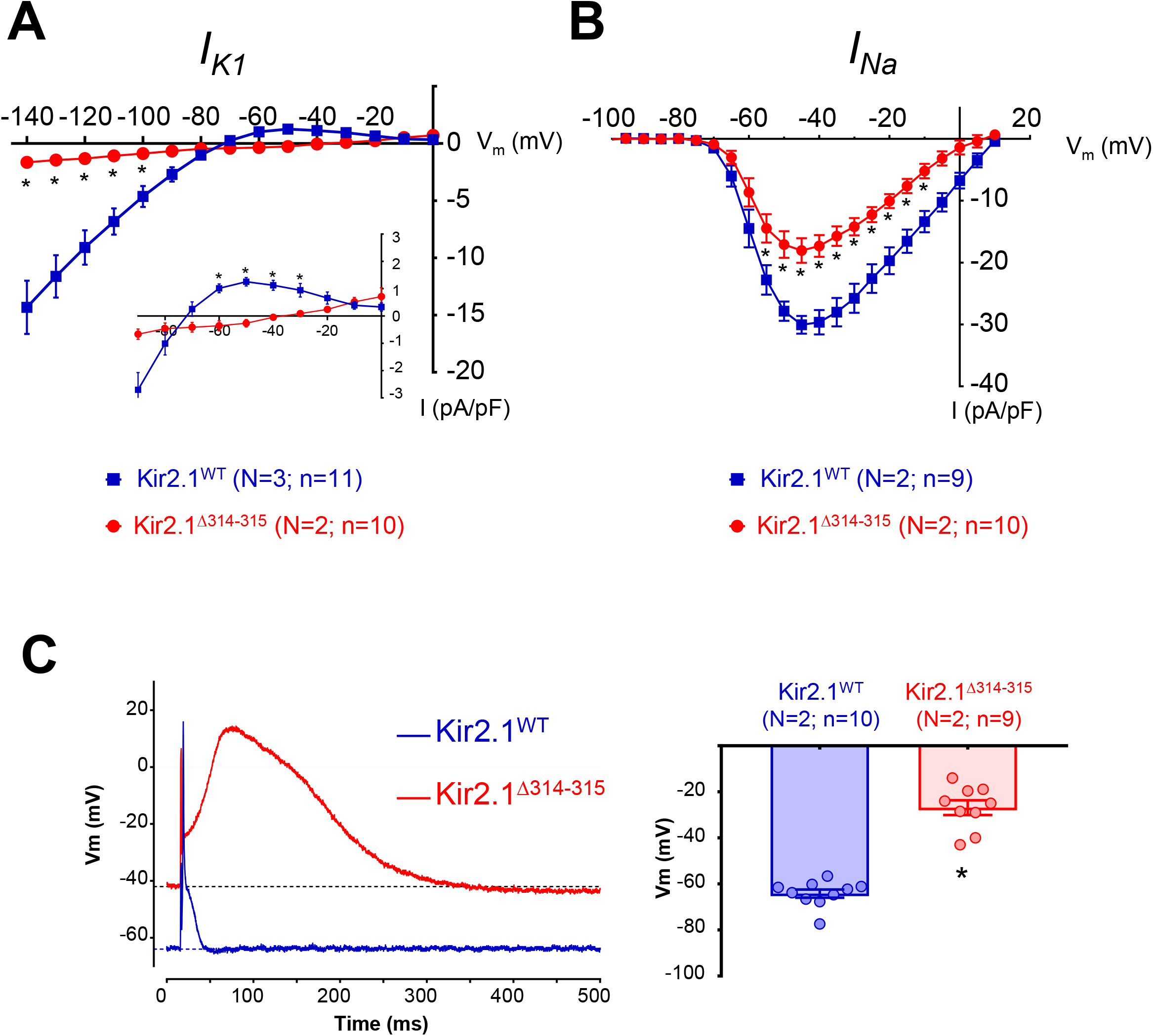
Cardiac expression of AAV-Kir2.1^Δ314-315^ results in significant reductions in *I_K1_* and *I_Na_* densities, depolarized resting membrane potential, reduced excitability and prolonged action potential duration. **A, B.** Current-voltage (IV) relationships of inward rectifying potassium current *I_K1_* (**A**) and inward sodium current *I_Na_* (**B**) in Kir2.1^WT^ and Kir2.1^Δ314-315^ ventricular cardiomyocytes. **C.** Representative action potential recordings obtained by current-clamping in Kir2.1^WT^ and Kir2.1^Δ314-315^ cardiomyocytes. Graph shows the quantification of resting membrane potential. Each value is represented as mean ± SEM. Statistical analyses were conducted using two-tailed t-test. *, p<0.05.

### Kir2.1 localizes in two defined microdomains in cardiomyocytes

Kir2.1 colocalizes with Nav1.5 at the cell membrane forming channelosomes in complex with various adaptor and scaffolding proteins at lateral membranes, t-tubules and intercalated discs structures^8, 10^. Representative confocal images of ventricular cardiomyocytes from control mice showed colocalization of Kir2.1 and Nav1.5 (**Figure 4A**), where RyR type-2 (RyR2) and SERCA also co-localize at the Z line (**Figure 4D**). Analysis of these confocal images also revealed that Kir2.1 localized in a defined structure running parallel to the t-tubule at ~0.9 μm (**Figure 4B-C**), halfway between two Z lines. The location of Kir2.1 at this microdomain corresponded to the M line where Kir2.1 colocalized with Ankyrin-B (**Figure 4E**). Furthermore, while a double Kir2.1 band pattern has not been specifically reported previously, it is clearly visible in several published illustrations from different laboratories ^10, 12, 17, 18^ (**Online Figure IV**). To test whether the double Kir2.1 band pattern was not mouse-specific we analyzed Kir2.1 immunolocalization in rat cardiomyocytes. Confocal imaging confirmed that Kir2.1 is localized in two distinct microdomains in both species (**Figure 4F**). More importantly, we also proved the presence of the double band pattern of Kir2.1 in mouse skeletal muscle tissue sections (**Figure 4G**), suggesting a potentially important role of both pools of Kir2.1 protein in different types of muscle cells.

**Figure 4.**
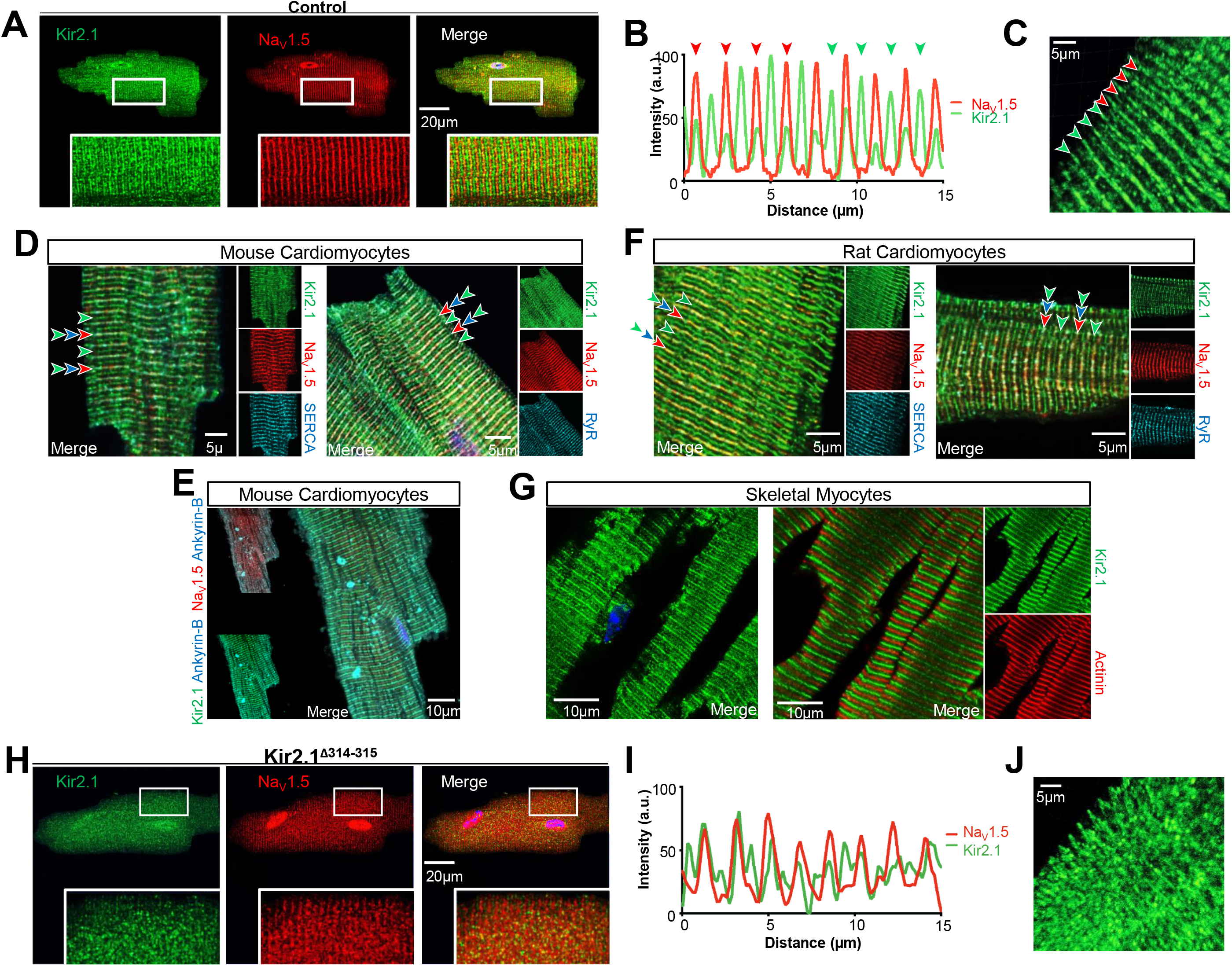
Immunostaining in cardiac and skeletal myocytes reveals two Kir2.1 bands at distinctly separate locations. Confocal images (**A**) and fluorescence intensity profiles (**B**) of isolated cardiomyocytes from control mice show the double Kir2.1 expression pattern in comparison with NaV1.5 channels. Arrows point to NaV1.5 channels in red and predominant Kir2.1 channels in green. Note the clearly discernible Kir2.1 expression pattern occurring at regular ~0.9 μm intervals while NaV1.5 appears at 1.8 μm intervals. **C.** Tridimensional reconstruction of Kir2.1 staining of bands close to NaV1.5 channels (red arrows) and those of separate Kir2.1 channels (green arrows). Confocal mouse (**D**) and rat (**F**) cardiomyocyte images showing the location of Kir2.1 channels together with SERCA (left) and RyR2 (right). **E.** Confocal images of isolated mouse cardiomyocyte from control mice showing the Kir2.1 expression pattern in comparison with NaV1.5 and Ankyrin-B. **G.** Confocal images of mouse skeletal muscle slices showing the double Kir2.1 localization pattern; Kir2.1 alone (left) and Kir2.1 plus actinin (right). **H, I.** Confocal images (**H**) and fluorescence intensity profiles (**I**) of isolated cardiomyocytes from Kir2.1^Δ314-315^ mice showing disrupted expression patterns of Kir2.1 and NaV1.5 channels. **J.** Tridimensional reconstitution of Kir2.1 staining showing the disorganization of the Kir2.1 expression pattern of a Kir2.1^Δ314-315^ cardiomyocyte.

Trafficking of both Kir2.1 and NaV1.5 to their membrane microdomains depends on their incorporation into clathrin-coated vesicles at the trans-Golgi network marked by interaction with the AP1 (adaptor protein complex 1) ϒ-adaptin subunit in both mice and rats ^10, 15^. As shown in **Figure 4H-J**, in contrast with the control group, Kir2.1^Δ314-315^ expression disrupts the well demarcated organization of both the Kir2.1-NaV1.5 channelosome band and the parallel band where Kir2.1 is expressed alone. On the other hand, as illustrated in **Online Figure V**, in WT mouse cardiomyocytes, AP1 ϒ-adaptin displays a clear colocalization with the NaV1.5 channelosome and little or no staining near the M line. However, the distribution is less organized in ATS1 mouse cardiomyocytes, showing a patchy distribution at a number of locations, likely due to the lack of a recognition site for interaction with AP1 in the Kir2.1^Δ314-315^ protein, a key mediator of Kir2.1 and NaV1.5 trafficking and membrane targeting ^10, 15^.

### A Kir2.1 protein subpopulation is localized at the SR membrane

To discriminate whether the new Kir2.1 protein band is in the sarcolemma or in an intracellular membrane compartment, we performed formamide-mediated detubulation assays (**Figure 5A-B**). Confocal images of cardiomyocytes stained to detect both Kir2.1 and NaV1.5 showed two clearly separate, alternating Kir2.1 protein localizations, one isolated and the other colocalizing with NaV1.5 channels (**Figure 5A**). However, upon formamide-mediated detubulation, the sarcolemmal membrane stained with wheat germ agglutinin (WGA) reflected t-tubular system disruption, and Kir2.1 and NaV1.5 staining was clearly obliterated (**Figure 5B**). In contrast, the unique Kir2.1 second structure associated with the M line remained intact despite detubulation, indicating its location in an intracellular compartment. Concordant with the data shown above and those described previously^8, 10, 19^, membrane fractionation using iodixanol-mediated density gradient showed that NaV1.5 channels segregated into two different populations, one Kir2.1-independent at the 10% fraction, and the other Kir2.1-dependent at the 15% and 20% fractions (**Figure 5C**). On the other hand, the largest proportion of Kir2.1 channels were NaV1.5-independent and isolated along with the specific SR protein calnexin (23% fraction), supporting the localization of Kir2.1 at the SR cellular domain (**Figure 4** and **Figure 5A-B**). Further, direct visualization of SR vesicles isolated together with cell nuclei revealed that Kir2.1 protein colocalized with SERCA, the major calcium transporter within the SR (**Figure 5D**). Altogether, the foregoing data demonstrate that the M line-associated Kir2.1 band is located at an SR microdomain.

**Figure 5.**
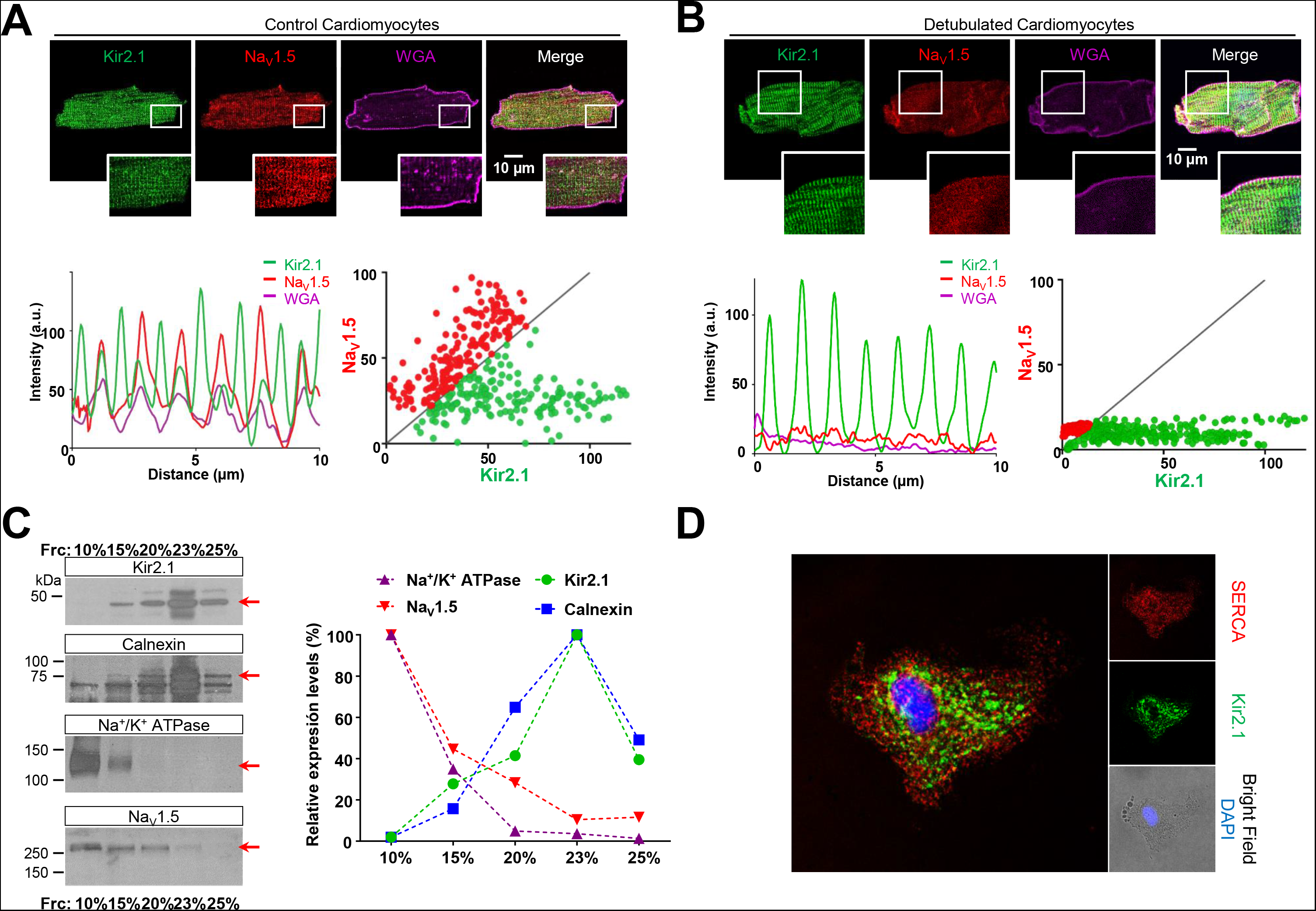
The new Kir2.1 band is intracellular and present at the SR membrane. **A. Top,** Confocal images of Kir2.1, NaV1.5 and cardiomyocyte membrane (WGA); **Bottom**, fluorescence profiles (right) and NaV1.5-Kir2.1 fluorescence correlation in isolated control (left). **B. Top,** Confocal images of Kir2.1, NaV1.5 and cardiomyocyte membrane (WGA) after formamide-mediated detubulation of cardiomyocytes reveals the intracellular Kir2.1 band alone. **Bottom**, fluorescence profiles show absence of Nav1.5 staining at the t-tubules (right), with absence of NaV1.5-Kir2.1 fluorescence correlation (left). **C. Left**, Kir2.1, Calnexin, Na^+^/K^+^ ATPase, and NaV1.5 analysis by western-blot after whole heart membrane fractionation in 10-25% of iodixanol. **Right,** Graph shows quantification of protein analysis shown in left. **D.** Confocal images of Kir2.1 and SERCA in the SR network connected with the envelope of an isolated nucleus.

### SR vesicular membranes display a rectifying SR K^+^ current sensitive to specific Kir2.1 channel modulators

To assess whether the SR Kir2.1 channels are functional, we carried out patch-clamp experiments in the SR vesicles around segregated nuclei (**Figure 6A**). Voltage-clamping revealed a magnesium-(**Figure 6B**) and spermine-sensitive (**Figure 6C-E**) potassium current with properties similar, but opposite in polarity to I_K1_ ^20, 21^ (**Figure 6C-E** and **Online Figure VI**), whose endogenous spermine-induced rectification was potentiated by exposure to asymmetrical potassium concentrations (**Figure 6F-G**) ^22^. Together, these data strongly suggest that SR Kir2.1 channels are functional and oriented to provide a K^+^-current that rectifies (decreases) in the direction of the SR lumen (hereafter termed “rectifying SR K^+^ current”).

**Figure 6.**
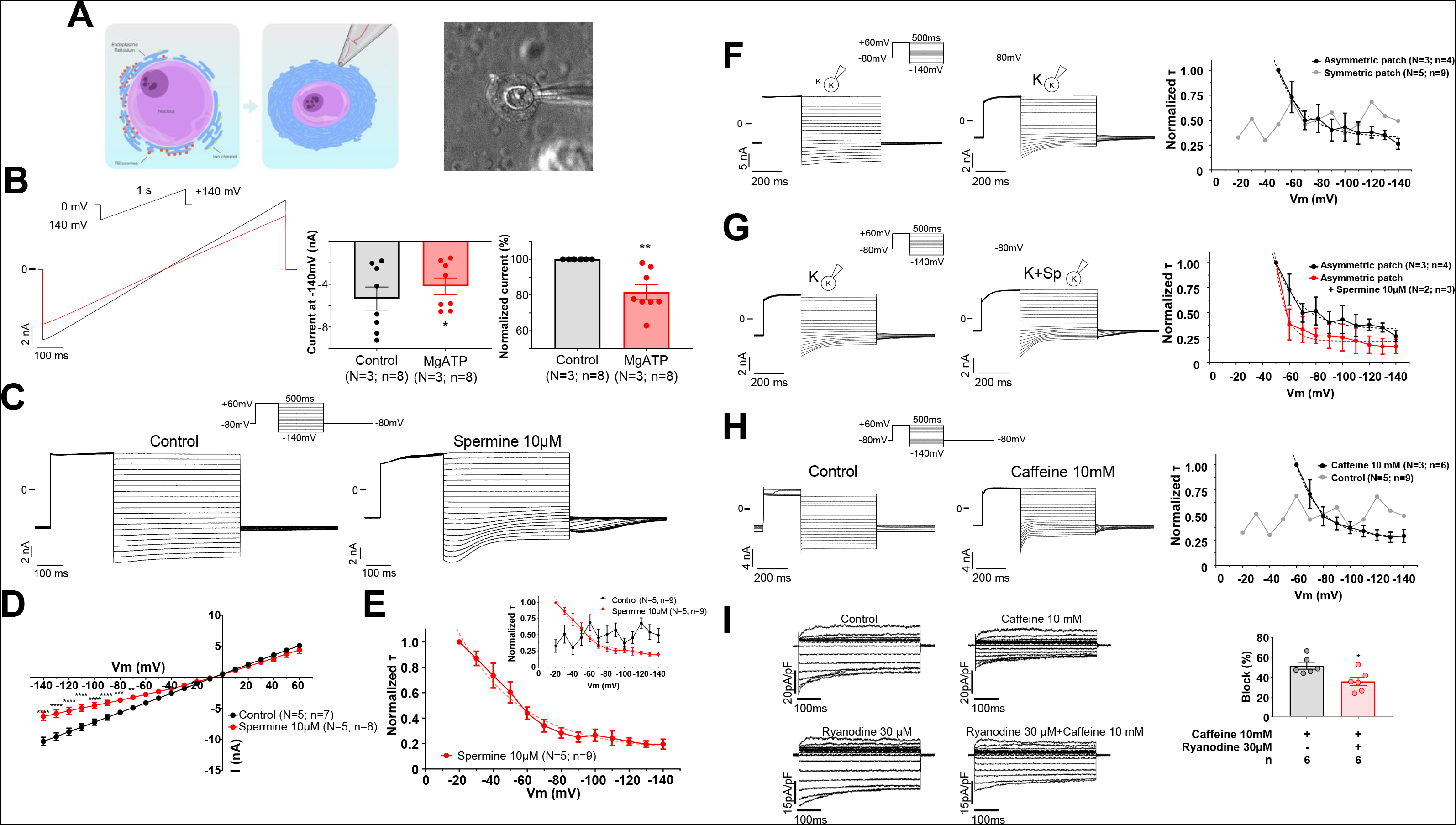
SR Kir2.1 is functionally active and shows inward-going rectification. **A.** Schematic representation (*left*) and phase contrast micrograph (*right*) of an isolated SR vesicle attached to the nucleus of a cardiomyocyte in a patch-clamp experiment. **B.** Representative patch-clamp recordings obtained by applying the voltage-ramp protocol shown on top in the absence (*black*) and the presence of 2mM MgATP. Graphs show the magnitude currents in absolute values (left) and normalized to each control experiment (right). **C.** Representative patch-clamp recordings obtained by applying the voltage protocol shown on top in the absence (*left*) and the presence (right) of spermine 10μM. **D.** Current-voltage (I-V) relationships constructed for the end of the test pulses in B for control (black) and spermine (red). **E.** Time constant (τ) of block by spermine 10μM (obtained from A) saturates at negative membrane potentials. Inset shows the same spermine data (red) compared to control (black). τ values were estimated using monoexponential fits. **F-H. Left**, representative patch-clamp experiment recordings using the voltage protocol shown on top. **Right**, time constant (τ) of block shown in left panels. **F.** Effect of residual endogenous spermine present in the SR vesicles under voltage-clamp conditions with symmetrical versus asymmetrical K^+^ concentration. **G.** Effect of spermine 10μM after asymmetrical patch-clamping. **H.** Effect of caffeine 10mM. τ values were estimated using monoexponential fits. **I. Left**, Representative recordings from HEK293 cells transfected with Kir2.1 channels before and after perfusion with caffeine 10mM. **Right,** Quantification of caffeine-induced block at −120mV. Each value is represented as mean ± s.e.m. Statistical analyses were conducted using two-tailed t-test and Two-way ANOVA test. *, p<0.05; **, p<0.01; ***, p<0.001; ****, p<0.0001.

The location and orientation of the rectifying SR Kir2.1 channels leads us to surmise that they contribute a countercurrent in the regulation of calcium movements across the SR, as has been suggested for K^+^ for many years ^23–25^. Thus, we analyzed the effect of caffeine perfusion, a strong RyR agonist, on the rectifying SR K^+^ current of the nuclear vesicles, which produced a strong spermine-like effect (**Figure 6H**). To determine whether caffeine acts directly on the Kir2.1 channel, or whether the effect is secondary to Ca^2+^ dynamic activation, we conducted similar experiments in HEK293 cells, in the absence and in the presence of 30μM ryanodine. The results demonstrated a direct effect of caffeine on the rectifying SR K^+^ current (**Figure 6I**). Altogether these data support the hypothesis that caffeine sensitive SR Kir2.1 channels have an important role in the control of the intracellular calcium dynamics, and that caffeine promotes Ca^2+^ efflux by acting simultaneously both as an agonist to RyR and as an antagonist to the rectifying SR K^+^ countercurrent.

### Calcium transient dynamics are defective in the ATS1 mutant mouse

The location and modulation of the new Kir2.1 band at the SR membrane leads us to speculate that this subgroup of channels likely plays a role in the control of intracellular calcium homeostasis. After all, the existence in the SR of potassium and other monovalent channels contributing to the bidirectional SR movement of calcium via RyR2 and SERCA has been suggested for many years ^26–29^. Thus, we analyzed the intracellular calcium dynamics in both, WT and ATS1 mice (**Figure 7**). Cardiomyocytes expressing Kir2.1^Δ314-315^ showed an excitation-contraction coupling defect in the form of slower calcium transient decay than WT cells both after field stimulation (**Figure 7A**) and caffeine administration (**Figure 7B**). Consequently, Kir2.1^Δ314-315^ cardiomyocytes showed multiple abnormal spontaneous calcium release events during systole and during diastole (**Figure 7A**). Altogether, our results provide a potential mechanism for the spontaneous and induced arrhythmias observed in the ATS1 mouse and to the phenotypic overlap between ATS1 and CPVT in some patients ^5, 6^.

**Figure 7.**
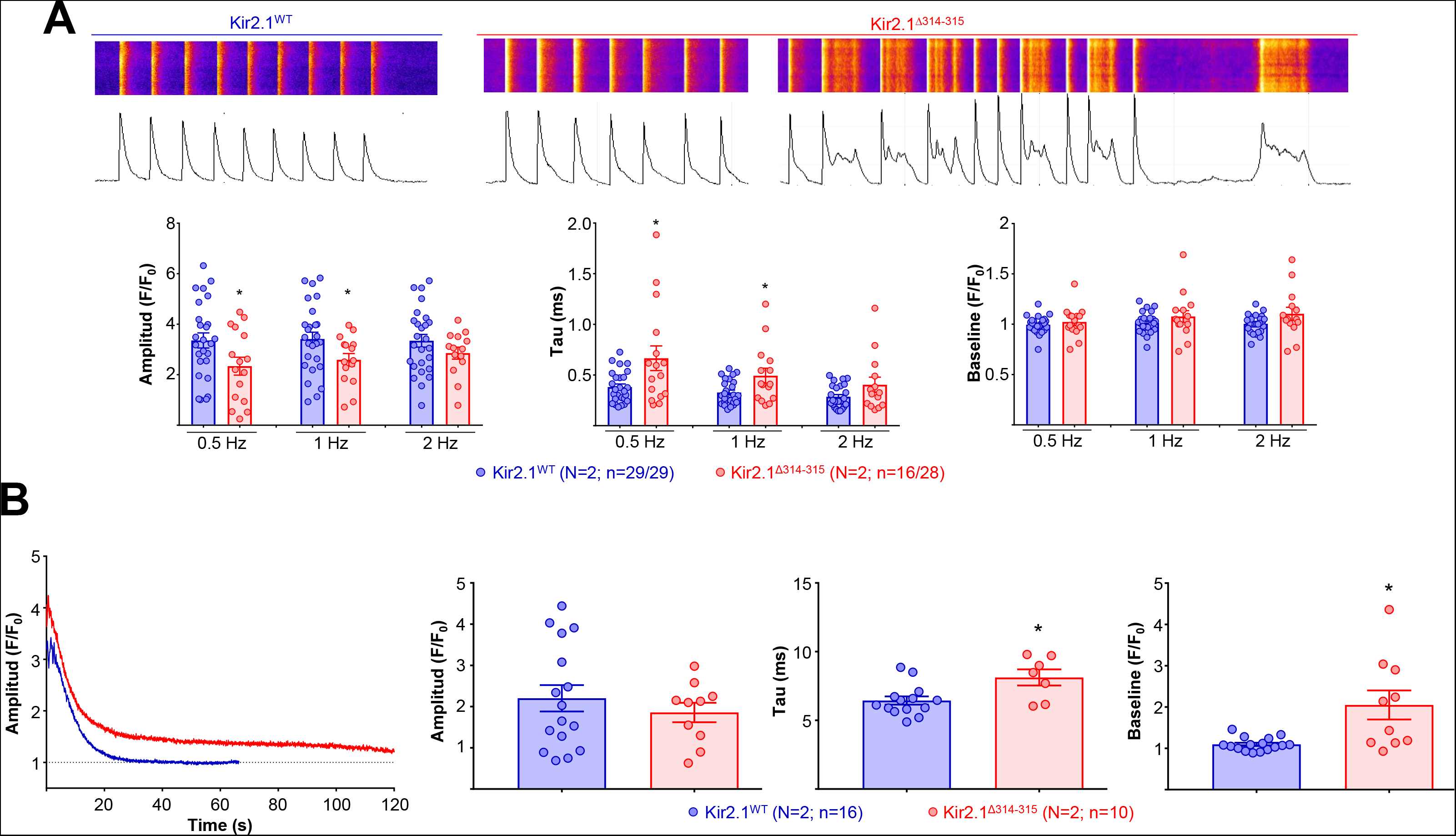
Calcium transient dynamics are defective in the ATS1 mutant mouse. **A.** Analysis of calcium dynamics in response to stimulation at 0.5, 1 and 2 Hz. Graphs show Amplitude, Tau (Decay kinetics) and Baseline of each Ca^2+^ transient. Note that in the representative Kir2.1^Δ314-315^ cardiomyocyte excitation-contraction coupling shows multiple abnormal spontaneous calcium release events during both systole and diastole, which are absent in the Kir2.1^WT^ cardiomyocyte. **B.** Analysis of Ca^2+^ transient decay after perfusion of caffeine 10mM during 20s.

## DISCUSSION

Using an AAV-mediated cardiac–specific mouse model of ATS1 expressing the trafficking deficient mutant Kir2.1^Δ314-315^ protein, we recapitulated the electrophysiological phenotype of ATS1 patients. In this disease model, arrhythmogenicity may be likely attributed to a dual dysfunction of the mutant Kir2.1 channels, one at the sarcolemma, resulting in reduced excitability and abnormal conduction, and the other at the SR membrane, where the mutant SR Kir2.1 channels directly alter intracellular calcium dynamics.

Our results demonstrate the existence of two spatially and functionally separate pools of Kir2.1 protein. The fist population colocalizes with NaV1.5 and AP1 near the Z band, and the other colocalizes with Ankyrin-B near the M line. Although *in vitro* experiments had already shown evidence of the formation of Kir2.1-NaV1.5 channelosomes at the cellular membrane^8, 10^, we are unaware of any previous studies reporting Kir2.1 localization at the M line microdomain. Using a combination of imaging, biochemical and electrophysiological approaches, we demonstrate herein that the latter pool of Kir2.1 channels is localized with and inverted polarity at the SR membrane. These distinct Kir2.1 protein microdomains are present in different species and different muscular cells, which suggests a conserved and generalized function.

A substantial amount of literature since the 1960’s has strongly suggested the existence of monovalent ion channels contributing countercurrent to the movements of calcium mediated by RyR2 and SERCA^26–29^. Such countercurrents are necessary to balance the movement of charges across the SR of both skeletal and cardiac muscle,^23^ in such a way that the Ca^2+^ equilibrium potential would be reached rapidly preventing any further release^25^. While calcium movement is accompanied by a voltage change across the SR membrane^24^, the RyR2– mediated SR movement of Ca^2+^ is well balanced by concomitant monovalent ion countercurrents^24^. To date, the identity of the monovalent channels at the SR had remained controversial ^23^, ^25^, ^30^, ^31^. Our calcium transient experiments in Kir2.1^Δ314-315^ cardiomyocytes suggest that SR Kir2.1 channel function may be important during diastole. If oriented as suggested by our voltage clamp experiments in nuclear vesicles (see **Figure 6**), SR Kir2.1 function could explain a fundamental current activity (from the SR lumen to the cytoplasm) giving rise to the countercurrent for SERCA-mediated Ca^2+^ reuptake, and to a lesser extent, RyR mediated Ca^2+^ release. In Kir2.1^Δ314-315^ cardiomyocytes, Kir2.1 was mislocalized and the two microdomains showed substantial disorganization. In these conditions, calcium transient dynamics was altered, as revealed by both prolonged recovery times of the calcium transients and multiple abnormal spontaneous calcium release events. Therefore, data presented here lead us to propose a potential molecular mechanism to explain the spontaneous arrhythmias as well as muscle weakness and periodic paralysis reported for ATS1 patients^5, 6^. We hypothesize that, in addition to disrupting the Kir2.1-NaV1.5 channelosome at the sarcolemma ^8, 10^, ATS1 also leads to dysfunction of SR Kir2.1 channels, directly altering SR countercurrents, thus disrupting e-c coupling and intracellular Ca^2+^ dynamics. This mechanism helps to understand the hitherto unexplained phenotypic overlap between ATS1 and CPVT in some patients, and the arrhythmias and intermittent paralysis seen in skeletal muscle diseases like hypokalemic periodic paralysis, hyperkalemic periodic paralysis, thyrotoxic periodic paralysis ^32, 33^ and malignant hyperthermia ^34^.

In accordance with previous reports ^1^, ^4^, ^35^, we demonstrate how *in-vivo* trafficking deficiency of one channel component of the Nav1.5-Kir2.1 macromolecular complex (in this case Kir2.1^Δ314–315^) negatively influences the functional localization of the other channel (Nav1.5), by disturbing channel trafficking to the plasma membrane. However, our data support the hypothesis that the stoichiometry of Kir2.1-Nav1.5 interaction at the membrane is unlikely to be 1:1, since voltage-clamp experiments show that more than 85% reduction of I_K1_ density at the membrane caused by cardiac Kir2.1^Δ314–315^ expression is accompanied by only ~40% reduction in *I_Na_* density. Such data highlight the idea that in mice, like rats and hiPSC-CMs, only a fraction of the Nav1.5 channels present in a cardiomyocyte share the AP-1 mediated Kir2.1 trafficking pathway as they reach the sarcolemma. Therefore, the data support the idea that Nav1.5 channels reach the plasma membrane by using more than one alternative pathways ^10^, ^19^.

As previous *in-vitro* experiments suggest ^15^, expression of Kir2.1 trafficking deficient mutant channels does not alter the global structure of the mutant monomere, which likely results in the formation of heterotetrameric Kir2.1^WT^-Kir2.1^Δ314–315^ channels translating into a dominant negative effect. As a consequence, decreases in both *I_K1_* and *I_Na_* upon Kir2.1^Δ314–315^ expression are due to alterations in the interaction of Kir2.1 and Nav1.5 with common protein partners ^8–10, 12, 36–40^. Here we show that the distribution of one of this partners, i.e., the AP1 protein, is compromised by the Kir2.1^Δ314–315^ mutation in the ATS1 mouse due to the lack of an AP1 binding site at the C-terminal SY_315_ residue, with consequent accumulation of the protein at the SR. These results lead us to hypothesize that proper Kir2.1-AP1 assembly taking place in the Golgi is necessary for the proper AP1-Nav1.5 interaction and trafficking to the sarcolemma. Accordingly, Nav1.5 channels may be exported from the Golgi in a signal-dependent manner through an AP1 clathrin adaptor interaction, as suggested previously ^10^. As such, trafficking vesicles may carry varying compositions of cargo proteins.

The availability of the ATS1 mouse model with cardiac-specific expression of Kir2.1^Δ314–315^ opens new possibilities for the understanding of the molecular mechanisms underlying the disease in patients. However, to date mutations in *KCNJ2* are the only genetic abnormalities identified in ATS patients who also present variable cardiac manifestations, despite the fact that approximately 60% of these patients have KCNJ2 mutations ^1, 4, 5^.

Although structural and functional heart disease has been described in some ATS1 patients ^41^, the most common cardiac defects are ECG and rhythm disturbances, including the presence of U waves, mild QT prolongation and conduction abnormalities like first degree atrioventricular block and bifascicular block ^42^. The arrhythmia burden is usually high, but surprisingly patients are for the most part asymptomatic ^35, 41, 43^. Nevertheless, cardiac arrest has been documented and family history of SCD has been identified ^41, 42^. The new AAV-mediated ATS1 mouse model recapitulates many of the above electrical abnormalities, including the high arrhythmia burden and susceptibility to VT/VF.

Furthermore, our data reveal that conduction and repolarization alterations, including PR, QRS and QT prolongation, as well as conduction velocity reduction demonstrated on ECGs of mice expressing Kir2.1^Δ314–315^, are a direct result of reduction of the density of currents generated by both Kir2.1 and Nav1.5 channels. These results extend previously described data for isolated rat and human cardiomyocytes^10^. Consistent with these results, our patch-clamp experiments demonstrate the reduction in *I_K1_* and *I_Na_* densities and consequently membrane depolarization and action potential duration prolongation. Moreover, the mouse phenotype is associated with e-c coupling defects, abnormal spontaneous calcium release, U waves and spontaneous arrhythmias mimicking CPVT, and increased susceptibility to AF, as well as VT/VF induced by intracardiac stimulation.

Importantly, we demonstrate that treatment with flecainide, a drug that is used to treat arrhythmias in ATS patients^44–46^, substantially exacerbates the ATS1 phenotype and is pro-arrhythmic in the mouse model. Clearly, the trafficking-deficient Kir2.1 mutation also disturbs Nav1.5 trafficking, ultimately contributing to further reduce excitability and impulse conduction velocity, establishing the substrate for life threatening arrhythmias. This calls for substantial caution and revaluation of the widespread use of flecainide in ATS1 patients, particularly those carrying trafficking deficient Kir2.1 mutations.

As an additional value, this model has permitted us to unravel a potential molecular mechanism for the phenotypic overlap between ATS1 and CPVT in some patients ^5, 6^, as well as a new actor in such an essential physiological function as e-c coupling in striated muscles. Based on the evidence presented here, we postulate that in addition to reduced *I_K1_* and *I_Na_* ^8, 10^, some ATS1 mutations also lead to dysfunction of SR Kir2.1 channels, directly altering SR countercurrents, disrupting e-c coupling and intracellular Ca^2+^ dynamics that result in stress-increased calcium-mediated arrhythmias that mimic CPVT.

To date, *KCNJ2* mutations are the only genetic abnormalities identified in ATS1 patients. Yet it is important to note the lack of consistency in its genotype– phenotype relationship in these patients stated above ^4^. In addition, the cardiac manifestations of the syndrome are variable ^43^, and the response to drugs is also likely to be variable, depending on the specific mutation. This new role for Kir2.1 channels at the SR opens new possibilities for the understanding of the molecular mechanisms underlying arrhythmias produced by other *KCNJ2* mutations. Revealing such mechanisms should lead to novel, more effective targets in the treatment of disorders related to calcium dynamic alterations including ATS and CPVT.

## Online Methods

See Expanded Methods in the Supplementary Information.

### Data Availability

The data, analytic methods, and study materials will be made available to other researchers for purposes of reproducing the results or replicating the procedures (available at the authors’ laboratories).

### Study Approval

All experimental and other scientific procedures using animals conformed to EU Directive 2010/63EU and Recommendation 2007/526/EC, enforced in Spanish law under *Real Decreto 53/2013*. Animal protocols were approved by the local ethics committees and the Animal Protection Area of the Comunidad Autónoma de Madrid (PROEX 019/17 and PROEX 111.4/20).

### Mice

C57BL/6J mice, 20-25-week-old, were obtained from Charles River Laboratories. Mice were reared and housed in accordance with institutional guidelines and regulations. The mice had free access to food and water. Mouse cardiomyocyte isolation and characterization was done as described previously (see Supplementary Information for details) ^8, 10, 19, 47^.

### Adeno-associated virus (AAV) vector production and purification

AAV vectors were produced by the triple transfection method, using HEK 293A cells as described previously^48, 49^. AAV vector titer (vg per ml) was carried out by quantitative real-time PCR as described^50^. Known copy numbers (10^5^–10^8^) of the respective plasmid (pAAV-empty vector, pAAV-Kir2.1^Δ314-315^ and pAAV-Luc) carrying the appropriate complementary DNA were used to construct standard curves.

### AAV injection

Mice were anesthetized with 100μl ketamine (60 mg/kg), xylazine (20 mg/kg) and atropine (9mg/kg) via the intraperitoneal route. Once asleep, animals were placed on a heated pad at 37±0.5°C to prevent hypothermia. Thereafter, 3.5×10^10^ virus particles were inoculated through the femoral vein in a final volume of 50μL, taking care to prevent introduction of air bubbles. Animals were then sedated with buprenorphine (s.c., 0.1 mg/kg) and maintained on the heating pad until recovery.

### AAV-Mediated Gene Distribution

Corporal distribution of protein expression was assessed as described^16^. Briefly, in-vivo bioluminescence signal was determined in luciferase control mice confirming the cardiac expression in mice.

### Surface electrocardiographic (ECG) recording

Mice were anesthetized using isoflurane inhalation (0.8-1.0% volume in oxygen), and efficacy of the anesthesia was monitored by watching breathing speed. Four-lead surface ECGs were recorded for 5 minutes, from subcutaneous 23-gauge needle electrodes attached to each limb using the MP36R amplifier unit (BIOPAC Systems).

### Intracardiac recording

An octopolar catheter (Science) was inserted through the jugular vein and advanced into the right atrium (RA) and ventricle As described previously^51^. Atrial and ventricular arrhythmia inducibility was assessed by applying 12–18 stimulus bursts. Atrial fibrillation (AF) was defined as the occurrence of rapid and fragmented atrial electrograms (lack of P waves) with irregular AV-nodal conduction and ventricular rhythm. Ventricular Tachycardia/Ventricular Fibrillation (VT/VF) was defined as an excessively fast monomorphic/polymorphic ventricular rhythm independent of atrial excitation.

### Cardiac Magnetic Resonance (CMR) imaging and analysis

Animals were anaesthetized with isoflurane and monitored for core body temperature, cardiac rhythm and respiration rate using a CMR compatible monitoring system. *In vivo* cardiac images were acquired using an Agilent VNMRS DD1 7T MRI system (Santa Clara, California, USA). All CMR images were analyzed using dedicated software (Segment software v1.9 R3819; http://segment.heiberg.se)^52^.

### Cardiac cell isolation

#### Cardiomyocyte isolation

The procedure was adapted from Garcia-Prieto et al^47^. Briefly, after euthanasia, the mouse heart was cannulated through the ascending aorta, retrogradely perfused and enzymatically digested.

#### Cardiofibroblast isolation

After heart digestion, the non-cardiomyocyte fraction was collected and centrifuged at 1,000g for 5 min and cells were re-suspended in DMEM/F-12 medium to adherence-mediated cardiofibroblast isolation.

### Sarcoplasmic reticulum vesicles isolation

Nuclei from cardiomyocytes were isolated as previously described ^53^. Briefly, isolated cardiomyocytes were washed and resuspended in Nuclei Isolation Solution (NIS). Finally, cell suspension was resuspended with a 30G syringe 40 times in order to induce cell lysis and nuclei release.

### Cardiomyocyte detubulation

The detubulation procedure was adapted from Kawai *et al.* ^54^ and Brette *et al.* ^55^.

### Immunofluorescence

#### Isolated cardiomyocytes

Cells were stained as described previously ^56^. Additional details are presented in the Expanded Methods section of the Supplemental Material.

#### SR vesicles

The immunofluorescence protocol described above was performed in nuclear samples that were decanted on SuperFrost Plus microscope slides and dried at 37°C for 1 hour.

### Calcium dynamics assays

Cytosolic Ca^2+^ was monitored according to previously described protocols ^56–58^.

### Membrane fractionation and immunoblotting

Ventricles from five mice were extracted and homogenized in ice-cold homogenization medium. After lysis, protein extract was processed on an OptiPrep Density Gradient Medium (DGM) ranging from 10 to 30% of iodixanol after centrifugation (according to manufacter’s specifications).

### Patch-clamping in isolated cardiomyocytes

Whole-cell and nuclear-vesicle patch-clamp techniques and data analysis procedures were similar to those described previously^9–13^. Details are presented in the Expanded Methods section of the Supplemental Material. The external and internal solutions are described on **Table S1**^10^, ^59^.

### Statistical analyses

We used GraphPad Prism software version 7.0 and 8.0. In general, comparisons were made using Student’s t-test. Unless otherwise stated, we used one- or two-way ANOVA for comparison among more than two groups. Data are expressed as mean ± SEM, and differences are considered significant at p<0.05.

## Acknowledgments

We thank to Dr. Carlos Galán-Arriola for his help in the graphical illustration of the manuscript. The project leading to these results has received funding from “la Caixa” Banking Foundation under the project code HR18-00304”. Support also came from a Severo Ochoa CNIC Intramural Project (Expediente 12-2016 IGP); a grant from Fundación La Marato TV3: Ayudas a la investigación en enfermedades raras 2020 (LAMARATO-2020); a grant from Instituto de Salud Carlos III to JJ; and MCIU grants BFU2016-75144-R to JAB and SAF2016-79490-R to VA from Ministerio de Ciencia e Innovación (MCIN), with co-funding from the *European Regional Development Fund* (“A way to build Europe”). The CNIC is supported by Instituto de Salud Carlos III (ISCIII), MCIN, and Pro CNIC Foundation.

## Author contribution statement

AM, JJ and JAB co-designed the experiments. AM, AGG, AIMM, FMC, and LKG performed most of the experiments with the exception of the membrane fractionation and in-vivo immunoblotting experiments, which were conducted by NGQ and MRM. VA participated in discussions and read the manuscript. FBJ, MLVP and IMC provided technical support, discussions and revisions. JJ and JAB provided supervision, funding and revisions. AM and JJ co-wrote the manuscript. All of the authors discussed the results, commented on and approved the manuscript.

## Competing interest statement

<u>The authors declare no competing interests.</u>

## Notes

### Competing Interest Statement

The authors have declared no competing interest.

## REFERENCES

1. Plaster, N.M. et al. Mutations in Kir2.1 cause the developmental and episodic electrical phenotypes of Andersen’s syndrome. Cell 105, 511–519 (2001).

2. Bendahhou, S. et al. Defective potassium channel Kir2.1 trafficking underlies Andersen-Tawil syndrome. J Biol Chem 278, 51779–51785 (2003).

3. Tawil, R. et al. Andersen’s syndrome: potassium-sensitive periodic paralysis, ventricular ectopy, and dysmorphic features. Ann Neurol 35, 326–330 (1994).

4. Tristani-Firouzi, M. et al. Functional and clinical characterization of KCNJ2 mutations associated with LQT7 (Andersen syndrome). J Clin Invest 110, 381–388 (2002).

5. Kukla, P., Biernacka, E.K., Baranchuk, A., Jastrzebski, M. & Jagodzinska, M. Electrocardiogram in Andersen-Tawil syndrome. New electrocardiographic criteria for diagnosis of type-1 Andersen-Tawil syndrome. Curr Cardiol Rev 10, 222–228 (2014).

6. Tully, I. et al. Rarity and phenotypic heterogeneity provide challenges in the diagnosis of Andersen-Tawil syndrome: Two cases presenting with ECGs mimicking catecholaminergic polymorphic ventricular tachycardia (CPVT). Int J Cardiol 201, 473–475 (2015).

7. Noujaim, S.F. et al. Up-regulation of the inward rectifier K+ current (I K1) in the mouse heart accelerates and stabilizes rotors. J Physiol 578, 315–326 (2007).

8. Milstein, M.L. et al. Dynamic reciprocity of sodium and potassium channel expression in a macromolecular complex controls cardiac excitability and arrhythmia. Proc Natl Acad Sci U S A 109, E2134–2143 (2012).

9. Matamoros, M. et al. Nav1.5 N-terminal domain binding to alpha1-syntrophin increases membrane density of human Kir2.1, Kir2.2 and Nav1.5 channels. Cardiovasc Res 110, 279–290 (2016).

10. Ponce-Balbuena, D. et al. Cardiac Kir2.1 and NaV1.5 Channels Traffic Together to the Sarcolemma to Control Excitability. Circ Res 122, 1501–1516 (2018).

11. Perez-Hernandez, M. et al. Pitx2c increases in atrial myocytes from chronic atrial fibrillation patients enhancing IKs and decreasing ICa,L. Cardiovasc Res 109, 431–441 (2016).

12. Park, S.S. et al. Kir2.1 Interactome Mapping Uncovers PKP4 as a Modulator of the Kir2.1-Regulated Inward Rectifier Potassium Currents. Mol Cell Proteomics 19, 1436–1449 (2020).

13. Abriel, H., Rougier, J.S. & Jalife, J. Ion channel macromolecular complexes in cardiomyocytes: roles in sudden cardiac death. Circ Res 116, 1971–1988 (2015).

14. Utrilla, R.G. et al. Kir2.1-Nav1.5 Channel Complexes Are Differently Regulated than Kir2.1 and Nav1.5 Channels Alone. Front Physiol 8, 903 (2017).

15. Ma, D. et al. Golgi export of the Kir2.1 channel is driven by a trafficking signal located within its tertiary structure. Cell 145, 1102–1115 (2011).

16. Cruz, F.M. et al. Exercise triggers ARVC phenotype in mice expressing a disease-causing mutated version of human plakophilin-2. J Am Coll Cardiol 65, 1438–1450 (2015).

17. Sengupta, S., Rothenberg, K.E., Li, H., Hoffman, B.D. & Bursac, N. Altering integrin engagement regulates membrane localization of Kir2.1 channels. J Cell Sci 132 (2019).

18. Yang, D. et al. MicroRNA Biophysically Modulates Cardiac Action Potential by Direct Binding to Ion Channel. Circulation 143, 1597–1613 (2021).

19. Perez-Hernandez, M. et al. Brugada syndrome trafficking-defective Nav1.5 channels can trap cardiac Kir2.1/2.2 channels. JCI Insight 3 (2018).

20. Lopatin, A.N., Shantz, L.M., Mackintosh, C.A., Nichols, C.G. & Pegg, A.E. Modulation of potassium channels in the hearts of transgenic and mutant mice with altered polyamine biosynthesis. J Mol Cell Cardiol 32, 2007–2024 (2000).

21. Hibino, H. et al. Inwardly rectifying potassium channels: their structure, function, and physiological roles. Physiol Rev 90, 291–366 (2010).

22. Kucheryavykh, Y.V. et al. Polyamine permeation and rectification of Kir4.1 channels. Channels (Austin) 1, 172–178 (2007).

23. Zsolnay, V., Fill, M. & Gillespie, D. Sarcoplasmic Reticulum Ca(2+) Release Uses a Cascading Network of Intra-SR and Channel Countercurrents. Biophys J 114, 462–473 (2018).

24. Sanchez, C. et al. Tracking the sarcoplasmic reticulum membrane voltage in muscle with a FRET biosensor. J Gen Physiol 150, 1163–1177 (2018).

25. Melzer, W. No voltage change at skeletal muscle SR membrane during Ca(2+) release-just Mermaids on acid. J Gen Physiol 150, 1055–1058 (2018).

26. Takeshima, H., Venturi, E. & Sitsapesan, R. New and notable ion-channels in the sarcoplasmic/endoplasmic reticulum: do they support the process of intracellular Ca(2)(+) release? J Physiol 593, 3241–3251 (2015).

27. Fink, R.H. & Veigel, C. Calcium uptake and release modulated by counter-ion conductances in the sarcoplasmic reticulum of skeletal muscle. Acta Physiol Scand 156, 387–396 (1996).

28. Miller, C. Voltage-gated cation conductance channel from fragmented sarcoplasmic reticulum: steady-state electrical properties. J Membr Biol 40, 1–23 (1978).

29. Costantin, L.L. & Podolsky, R.J. Depolarization of the internal membrane system in the activation of frog skeletal muscle. J Gen Physiol 50, 1101–1124 (1967).

30. Gillespie, D. & Fill, M. Intracellular calcium release channels mediate their own countercurrent: the ryanodine receptor case study. Biophys J 95, 3706–3714 (2008).

31. Yazawa, M. et al. TRIC channels are essential for Ca2+ handling in intracellular stores. Nature 448, 78–82 (2007).

32. Statland, J.M. et al. Review of the Diagnosis and Treatment of Periodic Paralysis. Muscle Nerve 57, 522–530 (2018).

33. Tsai, I.H. & Su, Y.J. Thyrotoxic periodic paralysis with ventricular tachycardia. J Electrocardiol 54, 93–95 (2019).

34. Schneiderbanger, D., Johannsen, S., Roewer, N. & Schuster, F. Management of malignant hyperthermia: diagnosis and treatment. Ther Clin Risk Manag 10, 355–362 (2014).

35. Tristani-Firouzi, M. & Etheridge, S.P. Kir 2.1 channelopathies: the Andersen-Tawil syndrome. Pflugers Arch 460, 289–294 (2010).

36. Gillet, L. et al. Cardiac-specific ablation of synapse-associated protein SAP97 in mice decreases potassium currents but not sodium current. Heart Rhythm 12, 181–192 (2015).

37. Lowe, J.S. et al. Voltage-gated Nav channel targeting in the heart requires an ankyrin-G dependent cellular pathway. J Cell Biol 180, 173–186 (2008).

38. Makara, M.A. et al. Ankyrin-G coordinates intercalated disc signaling platform to regulate cardiac excitability in vivo. Circ Res 115, 929–938 (2014).

39. Petitprez, S. et al. SAP97 and dystrophin macromolecular complexes determine two pools of cardiac sodium channels Nav1.5 in cardiomyocytes. Circ Res 108, 294–304 (2011).

40. Shy, D. et al. PDZ domain-binding motif regulates cardiomyocyte compartment-specific NaV1.5 channel expression and function. Circulation 130, 147–160 (2014).

41. Wilde, A.A. Andersen-Tawil syndrome, scarier for the doctor than for the patient? Who, when, and how to treat. Europace 15, 1690–1692 (2013).

42. Delannoy, E. et al. Cardiac characteristics and long-term outcome in Andersen-Tawil syndrome patients related to KCNJ2 mutation. Europace 15, 1805–1811 (2013).

43. Zhang, L. et al. Electrocardiographic features in Andersen-Tawil syndrome patients with KCNJ2 mutations: characteristic T-U-wave patterns predict the KCNJ2 genotype. Circulation 111, 2720–2726 (2005).

44. Mazzanti, A. et al. Natural History and Risk Stratification in Andersen-Tawil Syndrome Type 1. J Am Coll Cardiol 75, 1772–1784 (2020).

45. Miyamoto, K. et al. Efficacy and safety of flecainide for ventricular arrhythmias in patients with Andersen-Tawil syndrome with KCNJ2 mutations. Heart Rhythm 12, 596–603 (2015).

46. Rujirachun, P., Junyavoraluk, A., Pithukpakorn, M., Suktitipat, B. & Winijkul, A. Successful treatment of arrhythmia with beta-blocker and flecainide combination in pregnant patients with Andersen-Tawil syndrome: A case report and literature review. Ann Noninvasive Electrocardiol, e12798 (2020).

47. Garcia-Prieto, J. et al. beta3 adrenergic receptor selective stimulation during ischemia/reperfusion improves cardiac function in translational models through inhibition of mPTP opening in cardiomyocytes. Basic Res Cardiol 109, 422 (2014).

48. Xiao, X., Li, J. & Samulski, R.J. Production of high-titer recombinant adeno-associated virus vectors in the absence of helper adenovirus. J Virol 72, 2224–2232 (1998).

49. Hauswirth, W.W., Lewin, A.S., Zolotukhin, S. & Muzyczka, N. Production and purification of recombinant adeno-associated virus. Methods Enzymol 316, 743–761 (2000).

50. Prasad, K.R. et al. Topoisomerase Inhibition Accelerates Gene Expression after Adeno-associated Virus-mediated Gene Transfer to the Mammalian Heart. Mol Ther 15, 764–771 (2007).

51. Bao, Y. et al. Scn2b Deletion in Mice Results in Ventricular and Atrial Arrhythmias. Circ Arrhythm Electrophysiol 9 (2016).

52. Heiberg, E. et al. Design and validation of Segment--freely available software for cardiovascular image analysis. BMC Med Imaging 10, 1 (2010).

53. Mak, D.O., Vais, H., Cheung, K.H. & Foskett, J.K. Isolating nuclei from cultured cells for patch-clamp electrophysiology of intracellular Ca(2+) channels. Cold Spring Harb Protoc 2013, 880–884 (2013).

54. Kawai, M., Hussain, M. & Orchard, C.H. Excitation-contraction coupling in rat ventricular myocytes after formamide-induced detubulation. Am J Physiol 277, H603–609 (1999).

55. Brette, F., Komukai, K. & Orchard, C.H. Validation of formamide as a detubulation agent in isolated rat cardiac cells. Am J Physiol Heart Circ Physiol 283, H1720–1728 (2002).

56. Macias, A. et al. Paclitaxel mitigates structural alterations and cardiac conduction system defects in a mouse model of Hutchinson-Gilford progeria syndrome. Cardiovasc Res (2021).

57. Semenov, I. et al. Excitation and injury of adult ventricular cardiomyocytes by nano-to millisecond electric shocks. Sci Rep 8, 8233 (2018).

58. Brette, F., Despa, S., Bers, D.M. & Orchard, C.H. Spatiotemporal characteristics of SR Ca(2+) uptake and release in detubulated rat ventricular myocytes. J Mol Cell Cardiol 39, 804–812 (2005).

59. Moreno, C. et al. Modulation of voltage-dependent and inward rectifier potassium channels by 15-epi-lipoxin-A4 in activated murine macrophages: implications in innate immunity. J Immunol 191, 6136–6146 (2013).

