## Supplemental Material for "Dual Dysfunction of Kir2.1 Underlies Conduction and Excitation-Contraction Coupling Defects Promoting Arrhythmias in a Mouse Model of Andersen-Tawil Syndrome Type 1"

**Short Title:** Dual Function of Kir2.1 and ATS1 arrhythmogenesis

### These authors contributed equally to this work

*Corresponding authors:
José Jalife, MD, PhD.
Cardiac Arrhythmia Laboratory
Centro Nacional de Investigaciones Cardiovasculares
Melchor Fernández Almagro 3, 28029 Madrid, Spain


Juan A. Bernal, PhD.
Inherited Cardiomyopathies Lab & Head of the Viral Vector Unit (ViVU)
Centro Nacional de Investigaciones Cardiovasculares
Melchor Fernández Almagro 3, 28029 Madrid, Spain


#### *EXPANDED METHODS*

**Mice.** Wild-type, 20-25-week-old C57BL/6J mice were obtained from Charles River Laboratories. Mice were reared and housed in accordance with institutional guidelines and regulations. The mice had free access to food and water. Mouse cardiomyocyte isolation and characterization was done as previously described [1-4].

**Adeno-associated virus (AAV) vector production and purification.** AAV vectors were all produced by the triple transfection method, using HEK 293A cells as previously described[1]. AAV plasmids were cloned and propagated in the Stbl3 *E. coli* strain (Life Technologies). Shuttle plasmids pAAV-empty vector, pAAV-Kir2.1$^{\Delta 314\text{-}315}$ and pAAV-Luc were derived from pAcTnT (a gift from Dr B.A. French) and packaged into AAV-9 capsids with the use of helper plasmids pAdDF6 (providing the three adenoviral helper genes) and pAAV2/9 (providing rep and cap viral genes), obtained from PennVector. Shuttle vectors were generated by direct cloning (GeneScript) of synthesized fragments from NheI-SalI into pAcTnT cut with the same restriction enzymes.

The AAV shuttle and helper plasmids were transfected into HEK 293A cells by calcium-phosphate co-precipitation. A total of 840µg plasmid DNA (mixed in an equimolar ratio) was used per Hyperflask (Corning) seeded with $1.2 \times 10^8$ cells the day before. Seventy-two hours after transfection, the cells were collected by centrifugation and the cell pellet was resuspended in TMS (50 mmol/L Tris HCl, 150 mmol/L NaCl, 2 mmol/L $MgCl_2$) on ice before digestion with DNase I and RNaseA (0.1 mg/mL each; Roche) at 37 °C for 60 minutes. Clarified supernatant containing the viral particles was obtained by iodixanol gradient centrifugation.[2]

Gradient fractions containing virus were concentrated using Amicon UltraCel columns (Millipore) and stored at -70ºC.

Determination of AAV vector titer (vg per ml) were carried out by quantitative real-time PCR as described.[3] Known copy numbers ($10^5$–$10^8$) of the respective plasmid (pAAV-empty vector, pAAV- Kir2.1$^{\Delta 314\text{-}315}$ and pAAV-Luc) carrying the appropriate complementary DNA were used to construct standard curves.

**AAV injection.** Mice were anesthetized with 100µl of ketamine (60 mg/kg), xylazine (20 mg/kg) and atropine (9mg/kg) via the intraperitoneal route. Once asleep, animals were located on a heated pad at 37±0.5ºC to prevent hypothermia. A ~4-mm incision was made in the skin to expose the right femoral vein. To increase vessel diameter and facilitate infusion, blood flow was interrupted with a cotton bud for a couple of seconds. Once the vein was dilated, and insulin syringe vessel was introduced into the vein and $3.5\times10^{10}$ virus particles were inoculated in a final volume of 50µL, taking care to prevent introduction of air bubbles. Animals were then sedated with buprenorphine (s.c., 0.1 mg/kg) and maintained on the heating pad until recovery. Paracetamol was administered orally for 1 week.

**AAV-Mediated Gene Distribution.** Corporal distribution of protein expression was examined as previously described.[4] Briefly, four weeks after injection, in-vivo bioluminescence signal was performed in luciferase control mice confirming the cardiac expression by similar experiments in ex-vivo hearts organs from these mice. Infection efficiency was quantified by epi-fluorescence of whole hearts and microscopic images of cardiac slices in mock-injected and AAV-transduced mice.

**Electrocardiographic (ECG) recording.** *Surface ECG.-* Mice were anaesthetized using isoflurane inhalation (0.8-1.0% volume in oxygen), and efficacy of the anesthesia was monitored by watching breathing speed. Four-lead surface ECGs were recorded, for a period of 5 minutes, from subcutaneous 23-gauge needle electrodes attached to each limb using the MP36R amplifier unit (BIOPAC Systems).

During offline analysis, lead II was used for QRS duration using AcqKnowledge 4.1 analysis software. A representative 30s segment of the recording was averaged to obtain the signal-averaged ECG. QRS duration (before and after flecainide administration, 20 mg/Kg; and isoprenaline, 100 mg/Kg) was measured as the time interval between the earliest moment of deviation from baseline and the moment when the S-wave returned to the isoelectric line. QT duration was measured when the recording returned to the isoelectric line after T-wave and corrected by Framingham equation [5].

*Intracardiac recording* –An octopolar catheter (Science) was inserted through the jugular vein and advanced into the right atrium (RA) and ventricle as previously described.[7] Programmed electrical stimulation was assessed to determine the basal sinus node recovery time (SNRT), atrial refractory period (AERP), atrio-ventricular nodal effective refractory period (AVERP), and ventricular refractory period (VERP). Atrial and ventricular arrhythmia inducibility, was assessed by application of 12–18 atrial bursts and defined as the occurrence of rapid and fragmented electrograms (lack of regular P waves) with irregular AV-nodal conduction and ventricular rhythm.

**Cardiac Magnetic Resonance (CMR) imaging and analysis.** During CMR evaluation, animals were anaesthetized with isoflurane and were monitored for core body temperature, cardiac rhythm and respiration rate using a CMR compatible monitoring system. In vivo cardiac images were acquired using an Agilent VNMRS DD1 7T MRI system (Santa Clara, California, USA). A k-space segmented ECG-triggered cine gradient-echo sequence was used. After shimming optimization, cardiac four-chamber and left two-chamber views were acquired and used to plan the short axis sequence. Mice were imaged with the following parameter settings: number of slices, 13; slice thickness, 0.8 mm; gap, 0 mm; matrix size, 256x256; field of view, 30x30 mm2; gating: ECG and respiratory triggered; cardiac phases, 20; averages, 4; effectiveTE, ~1.8 ms, minimumTR, 7 ms; flip angle, 25º; trigger delay, 2 ms; trigger window, 8 ms; dummy scans, 2.

All CMR images were analyzed using dedicated software (Segment software v1.9 R3819; http://segment.heiberg.se).[8] Images were analyzed by two experienced observers blinded to the study allocation. All CMR images were of good quality and could be analyzed. The short-axis data set was analyzed quantitatively by manual detection of endocardial borders in end-diastole and end-systole, with exclusion of papillary muscles and trabeculae, in order to obtain both left and right end-diastolic volume, end-systolic volume and ejection fraction.

**Cardiac echocardiography.** Transthoracic echocardiography was blindly performed by an expert operator using a high-frequency ultrasound system (Vevo 2100, Visualsonics Inc., Canada) with a 40-MHz linear probe. Two-dimensional (2D) and M-mode (MM) echography were performed at a frame rate above 230 frames/sec, and pulse wave Doppler (PW) was acquired with a pulse repetition frequency of 40 kHz. Mice were lightly anesthetized with 0.5-2% isoflurane in oxygen, adjusting the isoflurane delivery trying to maintain the heart rate in

450±50 bpm. Mice were placed in supine position using a heating platform and warmed ultrasound gel was used to maintain normothermia. A base apex ECG was continuously monitored. Images were transferred to a computer and were analysed off-line using the Vevo 2100 Workstation software. For left ventricular (LV) systolic function assessment, parasternal standard 2D and MM, long and short axis views (LAX and SAX view, respectively) were acquired. LV ejection fraction and chamber dimensions were calculated from these views.

**Cardiac cell isolation.** *Cardiomyocyte isolation*.- The procedure was adapted from Garcia-Prieto et al.[4] Briefly, after euthanasia in a $CO_2$ chamber, mice were placed in the supine position, and the ventral thoracic region was wiped with 70% alcohol. The heart was quickly removed and incubated at room temperature (RT) in $Ca^{2+}$-free Perfusion-Buffer (PB; in mmol/L): NaCl, 113; KCl, 4.7; $KH_2PO_4$, 0.6; $Na_2HPO_4$, 0.6; $MgSO_4{\cdot}7H_2O$, 1.2; $NaHCO_3$, 12; $KHCO_3$, 10; Phenol Red, 0.032; HEPES, 0.922; taurine, 30; glucose, 5.5; 2,3-butanedione-monoxime, 10; pH 7.4. Fat was cleaned and the heart was cannulated through the ascending aorta and mounted on a modified Langendorff-perfusion apparatus. The heart was then retrogradely perfused (1 mL/min) for 5 min at 37°C with PB. Enzymatic digestion was performed with digestion-buffer (DB): PB supplemented with Liberase™ (0.2 mg/mL), Trypsin 2.5 % (5.5 mmol/L); DNase ($5x10^{-3}$ U/mL) and $CaCl_2$ (12.5 µM)] for 20 min at 37°C. At the end of enzymatic digestion, both ventricles were isolated and gently disaggregated in 3 mL DB. The resulting cell suspension was filtered through a 200-µm sterile mesh (SEFAR-Nitex) and transferred for enzymatic inactivation to a tube with 12 mL of stopping-buffer-1 (SB-1): PB supplemented with fetal bovine serum (FBS, 10 % v/v) and $CaCl_2$ (12.5 µmol/L). After gravity sedimentation for 20 min, supernatant was removed, and cardiomyocytes were resuspended in stopping-buffer-2 (SB-2) containing lower FBS (5% v/v) for another 20 min. Cardiomyocyte $Ca^{2+}$-reintroduction was performed in SB-2 with two progressively increased $CaCl_2$ concentrations (0.112 and 1 mmol/L). Cells were resuspended and allowed to decant for 15 min in each step, contributing to the purification of the cardiomyocyte suspension.

*Cardio-fibroblast isolation.-* After SB-1 addition in the cardiomyocytes isolation protocol, the supernatant was collected and centrifuged at 1,000g for 5 min and cells were re-suspended in DMEM/F-12 medium. To perform adherence-mediated fibroblast isolation, cell suspension was seeded in a 24-well culture dish during 30-45min. After that, cells not adhered at the bottom of the dish were removed by gently washing 2-3 times with PBS. For image acquisition cells were stored in culture for 3 days changing the medium every day.

**Sarcoplasmic reticulum (SR) vesicles isolation.-** Nuclei from isolated cardiomyocytes were isolated as previously described [6]. Briefly, isolated cardiomyocytes were washed with ice-cold PBS, centrifuged and the cell pellet was resuspended in an appropriate volume of ice-cold Nuclei Isolation Solution (NIS; in mM: 150 KCl, 250 Sucrose, 10 Tris-HCl, 1.4 β-Mercaptoethanol, pH 7.3 KOH supplemented with one tablet of complete protease inhibitor cocktail for each 40 ml of NIS, Roche Applied Science 1697498). Usually, a whole heart was divided in 3 aliquots and the cells resuspended in 250µl of NIS. Finally, SR vesicles were obtained by resuspending the cell suspension with a 30G syringe 40 times in order to induce cell lysis and nuclei release.

**Cardiomyocyte detubulation.** The detubulation procedure was adapted from Kawai *et al.* [7] and Brette *et al.* [8] Briefly, isolated cardiomyocytes were washed with a bath solution (BS; in mM): 113 NaCl, 5 KCl, 1 $MgSO_4$, 1 $CaCl_2$, 1 $Na_2HPO_4$,

20 sodium acetate, 10 glucose, 10 HEPES and 5 U/L insulin, pH to 7.4 NaOH. Cell detubulation was performed after 15 min at room temperature in detubulation solution (DS): DS supplemented with formamide 1.5 M and EGTA 5.25 µM. After this, formamide was washed-out with BS.

**Immunofluorescence.** *Isolated cardiomyocytes.-* Cells were fixed in 4% formaldehyde in PBS at RT, shaken gently for 10 min, washed with PBS, and stored at 4ºC in PBS until use. Cells were then incubated for 10 min at RT with wheat germ agglutinin (WGA) Alexa488 (W11261, ThermoScientific, 1/100) under gentle shaking, washed with PBS, and re-fixed in 4% formaldehyde in PBS at RT with gentle shaking for 15 min to avoid dye internalization. Thereafter, cells were prepared for immunofluorescence. Cells were blocked and permeabilized for 90 min at RT in PBS containing 0.2% Triton X-100 and 10% normal goat serum, and incubated overnight at 4°C with anti-Kir2.1 (1:200, APC-026, Alomone Labs), anti-Nav1.5 (1:50, AGP-008, Alomone Labs), anti-SERCA (1:200, Santa Cruz Biotechnology), anti-RyR2 (1:200, ThermoFisher Scientific), anti-Ankyrin-B (1:200, Santa Cruz Biotechnology) and anti-Actinin (1:200, MERCK). Samples were then incubated for 2h at RT with secondary antibodies (1/500 in all cases) and mounted in Fluoroshield™ -DAPI imaging medium (F6057, Merck). Images of individual cardiomyocytes were acquired with a Leica SP8 confocal microscope. Finally, Imaris (Bitplane) software was used for 3D rendering.

*SR vesicles.-* The immunofluorescence protocol described above was performed in nuclear samples that were decanted on SuperFrost Plus microscope slides and dried at 37ºC for 1 hour.

**Calcium dynamics assays.** Cytosolic $Ca^{2+}$ was monitored as previously described[9, 10]. Briefly, cells were loaded with Fluo-4-AM (Invitrogen, Carlsbad, CA) by incubation for 15 min in Tyrode's solution containing 5 μM Fluo-4-AM and 0.02% Pluronic F-127 (Life Technologies, Grand Island, NY), in the dark at RT. Cells were allowed to settle to the bottom of the perfusion chamber (RC-26, Warner Instruments) mounted on the stage of an inverted LSM 880 Carl Zeiss confocal microscope before being perfused with the corresponding solution. All experiments were performed at RT. Images were taken using a 20X, NA 0.8 dry objective. Fluo-4-AM fluorescence was detected in line scan mode (usually 2 ms/scan), with the line drawn approximately through the center of the cell parallel to its long axis. Fluo-4-AM was excited with a blue laser (488 nm), and emission was detected between 505 and 605 nm.

**Membrane fractionation and immunoblotting.** Ventricles from five mice were extracted and homogenized in ice-cold homogenization medium (HM; 250 mM sucrose, 10mM Hepes-NaOH pH 7.4, 1mM EDTA, 1mM EGTA complemented with a mixture of protease inhibitor (Roche)) using a glass potter homogenizer and then passed through syringe with a 25G needle ten times. The total extract was centrifuged at 1,500g for 10 minutes at 4ºC to remove non-disrupted cells and the post-nuclear fraction. Supernatant was supplemented with 3 volumes of HM and centrifugated at 38,400g for 2 hours at 4ºC. The crude pellet was processed on an OptiPrep Density Gradient Medium (DGM) ranging from 10 to 30% of iodixanol prepared as described in the datasheet (Alere Technologies AS). After centrifugation at 130,000g overnight (16h) at 4ºC on a SW32Ti rotor, 4 fractions were isolated and further subjected to a 3h centrifugation at 170,000g at 4ºC on a SW40Ti rotor. Each final pellet was resuspended in RIPA buffer

($10mM\ PO_4Na_2/K$ buffer pH 7.2, 150mM NaCl, 1g/100ml sodium deoxycholate, 1% Triton X-100, 1% Nonidet P40) supplemented with a mixture of protease inhibitors (Roche). Identical volumes of each fraction were separated on 8 and 10% SDS-PAGE gels. Primary antibodies were rabbit anti-Kir 2.1 (APC-026, Alomone), mouse anti-calnexin (MA3-027, Invitrogen), mouse anti-ATPase (ab7671, Abcam) and rabbit anti-Nav1.5 (*SCN5A*, Alomone).

**Patch-clamping in isolated cardiomyocytes.** Whole-cell voltage and current-clamp recordings and data analysis procedures were similar to those previously described [9–13]. The external and internal solutions are described on **Online Table I.**

Aliquots of cardiomyocytes were placed in a superfusion chamber (RC-26, Warner Instruments) mounted on the stage of an inverted microscope (DMi8, Leica). Cells were allowed to settle on the bottom of the perfusion chamber before being perfused with the corresponding solution. After patch rupture, whole-cell voltage or current-clamp recordings were made using an Axopatch 200 B amplifier (Axon Instruments). Pipettes made from borosilicate glass (GD-1, Narishige, OD: 1 mm; ID: 0.6 mm) had resistances of 1-3 MΩ. Series resistance compensation of 80-90% was achieved. All voltage-clamp currents were low-pass filtered at 2 kHz with an analog filter, and digitized at (4-10 Hz). For $Ca^{2+}$ current experiments, cells were stabilized 5 minutes in the whole-cell configuration before starting the voltage-clamp protocols to control for current run-down. Data were recorded using pClamp 10.0 software with Clampex 10.0 program and analyzed with the Clampfit 10.0 program (Axon Instruments, Foster City, USA). Current amplitudes were normalized to the cell capacitance to account for differences in cell size, and expressed as densities (pA/pF).

*Action potential (AP) recordings.-* Threshold current was determined using one-msec pulses at increasing amplitudes (0.2 nA/pulse) and frequency of 1 Hz. Thereafter, AP were evoked by the injection of 1 msec pulses of constant amplitude. AP durations (APD) was measured at 20, 50, 70 and 90% of repolarization.

Current-voltage (*IV) relationships.- Potassium currents.* IV relationship for the inwardly rectifying K+ current ($I_{K1}$) were constructed from the current changes produced by a 500 ms voltage-clamp step applied in 10 mV increment from –110 to +50 mV from a holding potential of –80 mV at 0.1 Hz. $I_{K1}$ was calculated by subtracting currents recorded in the absence or presence of 500µM $BaCl_2$. We measured $I_{to}$ as the difference current, obtained by subtracting records with an inactivating pre-pulse voltage-clamp protocol (100 ms at –40mV) from those without the inactivating pre-pulse.

*Sodium currents*.- To record the sodium ($I_{Na}$) current-voltage relationship, cells were held at –160 mV and stepped for 100 ms from -100 mV up to +5 mV in 5 mV increments at 0.2 Hz. $I_{Na}$ was measured at the peak. In both cases leak currents were subtracted using the P/4 protocol.

*Isolated SR vesicles.-* After nuclei isolation, the sample was first stained with Sytox-green 5µM (S7020, Thermofisher) before experiments for clearer nuclei discrimination. Then, small aliquots were placed in a perfusion chamber (RC-26, Warner Instruments) mounted on the stage of an inverted microscope (DMi8, Leica). Samples were allowed to settle on the bottom of the perfusion chamber before being perfused with the corresponding solution. After patch rupture, whole-vesicle voltage-clamp recordings were made using an Axopatch 200B amplifier (Axon Instruments). Pipettes made from borosilicate glass (GD-1,

Narishige, OD: 1 mm; ID: 0.6 mm) had resistances of 6-8 MΩ. Series resistance compensation of 80-90% was achieved. All voltage-clamp currents were low-pass filtered at 2 kHz with an analog filter, and digitized at (4-10 Hz). Data were recorded using pClamp 10.0 software with Clampex 10.0 program and analyzed with the Clampfit 10.0 program (Axon Instruments, Foster City, USA). Whole-vesicle patch-clamping experiments were recorded by using $Mg^{2+}$- and spermine-free solution on both sides of the patch containing (in mM): 123 KCl, 5 $K_2EDTA$, 7.2 $K_2HPO_4$, and 8 $KH_2PO_4$ (pH 7.2 KOH).

*HEK293 cells.-* The external and internal solutions are described on **Online Table I.** IV relationship were constructed as indicated from the current changes produced by a 500 ms voltage-clamp step applied in 10 mV increment from –110 to +50 mV from a holding potential of –80 mV at 0.1 Hz, as previously reported [2, 11]. $I_{K1}$ was calculated by subtracting currents recorded in the absence or presence of 500µM $BaCl_2$.

**Statistical analyses**. Statistical analyses were performed using GraphPad Prism software version 7.0 and 8.0. Comparisons were generally made by Student's t-test. Unless otherwise stated, we used one- or two-way ANOVA for comparison between more than two groups. Data are expressed as mean ± s.e.m., and differences are considered significant at $p<0.05$.

**Online Table I: External and internal solutions used in patch-clamp experiments.**

| Product | $K^+$ Currents | | $Na^+$ Currents | |
|---|---|---|---|---|
| | Bath solution (mM) pH 7.4 (NaOH) | Internal solution (mM) pH 7.2 (KOH) | Bath solution (mM) pH 7.35 (CsOH) | Internal solution (mM) pH 7.2 (CsOH) |
| Aspartic acid | - | - | - | - |
| Calcium chloride ($CaCl_2$) | 1 | 1 | 1 | - |
| Cesium chloride (CsCl) | - | - | 132.5 | - |
| Cesium Fluoride (CsF) | - | - | - | 135 |
| Cesium Hydroxide (CsOH) | - | - | - | - |
| EGTA | - | 10 | - | 10 |
| Glucose | 5.5 | - | 10 | - |
| HEPES | 10 | 5 | 20 | 5 |
| K-Aspartate | - | 110 | - | - |
| $K_2ATP$ | - | 4 | - | - |
| Potassium Chloride (KCl) | 5.4 | 20 | - | - |
| Magnesium Adenosine-Tri-Phosphate (MgATP) | - | - | - | 5 |
| Magnesium Chloride ($MgCl_2$) | 1 | 1 | 1 | - |
| Niquel (II) Chloride ($NiCl_2$) | - | - | 1 | - |
| Sodium Chloride (NaCl) | 130 | 8 | 5 | 5 |
| $Na_2HPO_4$ | 0.33 | - | - | - |
| Tetraethylammonium (TEA) | - | - | - | - |

#### *REFERENCES*


1. Milstein, M.L. *et al.* Dynamic reciprocity of sodium and potassium channel expression in a macromolecular complex controls cardiac excitability and arrhythmia. *Proc Natl Acad Sci U S A* **109**, E2134-2143 (2012).
2. Ponce-Balbuena, D. *et al.* Cardiac Kir2.1 and NaV1.5 Channels Traffic Together to the Sarcolemma to Control Excitability. *Circ Res* **122**, 1501-1516 (2018).
3. Perez-Hernandez, M. *et al.* Brugada syndrome trafficking-defective Nav1.5 channels can trap cardiac Kir2.1/2.2 channels. *JCI Insight* **3** (2018).
4. Garcia-Prieto, J. *et al.* beta3 adrenergic receptor selective stimulation during ischemia/reperfusion improves cardiac function in translational models through inhibition of mPTP opening in cardiomyocytes. *Basic Res Cardiol* **109**, 422 (2014).
5. Sagie, A., Larson, M.G., Goldberg, R.J., Bengtson, J.R. & Levy, D. An improved method for adjusting the QT interval for heart rate (the Framingham Heart Study). *Am J Cardiol* **70**, 797-801 (1992).
6. Mak, D.O., Vais, H., Cheung, K.H. & Foskett, J.K. Isolating nuclei from cultured cells for patch-clamp electrophysiology of intracellular Ca(2+) channels. *Cold Spring Harb Protoc* **2013**, 880-884 (2013).
7. Kawai, M., Hussain, M. & Orchard, C.H. Excitation-contraction coupling in rat ventricular myocytes after formamide-induced detubulation. *Am J Physiol* **277**, H603-609 (1999).
8. Brette, F., Komukai, K. & Orchard, C.H. Validation of formamide as a detubulation agent in isolated rat cardiac cells. *Am J Physiol Heart Circ Physiol* **283**, H1720-1728 (2002).
9. Semenov, I. *et al.* Excitation and injury of adult ventricular cardiomyocytes by nano- to millisecond electric shocks. *Sci Rep* **8**, 8233 (2018).
10. Brette, F., Despa, S., Bers, D.M. & Orchard, C.H. Spatiotemporal characteristics of SR Ca(2+) uptake and release in detubulated rat ventricular myocytes. *J Mol Cell Cardiol* **39**, 804-812 (2005).
11. Moreno, C. *et al.* Modulation of voltage-dependent and inward rectifier potassium channels by 15-epi-lipoxin-A4 in activated murine macrophages: implications in innate immunity. *J Immunol* **191**, 6136-6146 (2013).
12. Sengupta, S., Rothenberg, K.E., Li, H., Hoffman, B.D. & Bursac, N. Altering integrin engagement regulates membrane localization of Kir2.1 channels. *J Cell Sci* **132** (2019).
13. Yang, D. *et al.* MicroRNA Biophysically Modulates Cardiac Action Potential by Direct Binding to Ion Channel. *Circulation* **143**, 1597-1613 (2021).
14. Park, S.S. *et al.* Kir2.1 Interactome Mapping Uncovers PKP4 as a Modulator of the Kir2.1-Regulated Inward Rectifier Potassium Currents. *Mol Cell Proteomics* **19**, 1436-1449 (2020).
15. Nichols, C.G., Makhina, E.N., Pearson, W.L., Sha, Q. & Lopatin, A.N. Inward rectification and implications for cardiac excitability. *Circ Res* **78**, 1-7 (1996).

***SUPPLEMENTAL FIGURES***

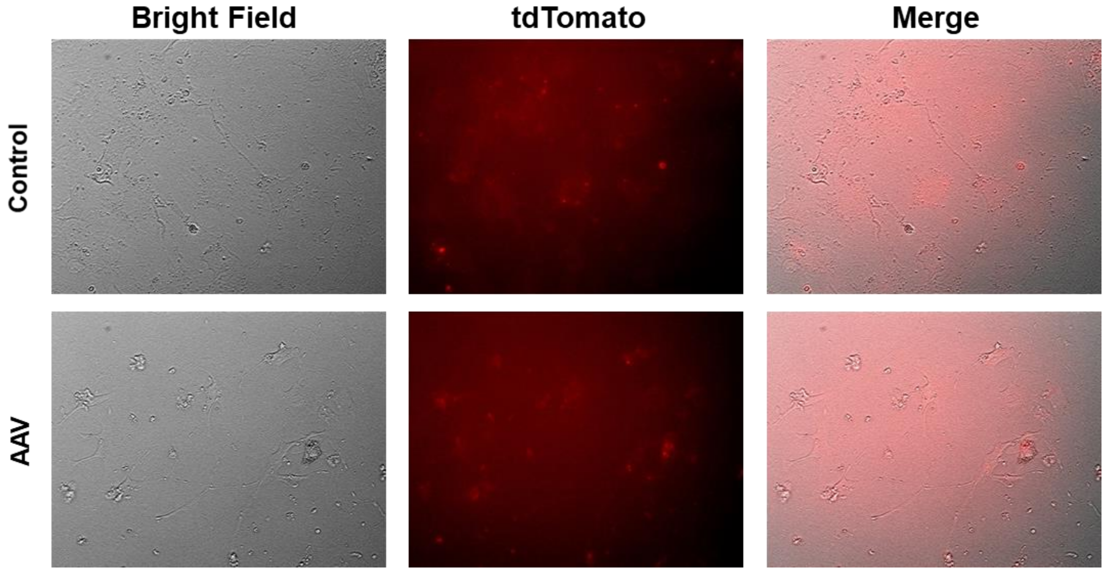


**Online Figure I. AAV-Kir2.1$^{\Delta 314\text{-}315}$ associated with the cTnT promoter is not expressed in cardiac fibroblasts.** Representative fluorescence images of cardio-fibroblast isolated from control and AAV-transduced mice to compare expression levels of tdTomato in non-cardiomyocytes cells.

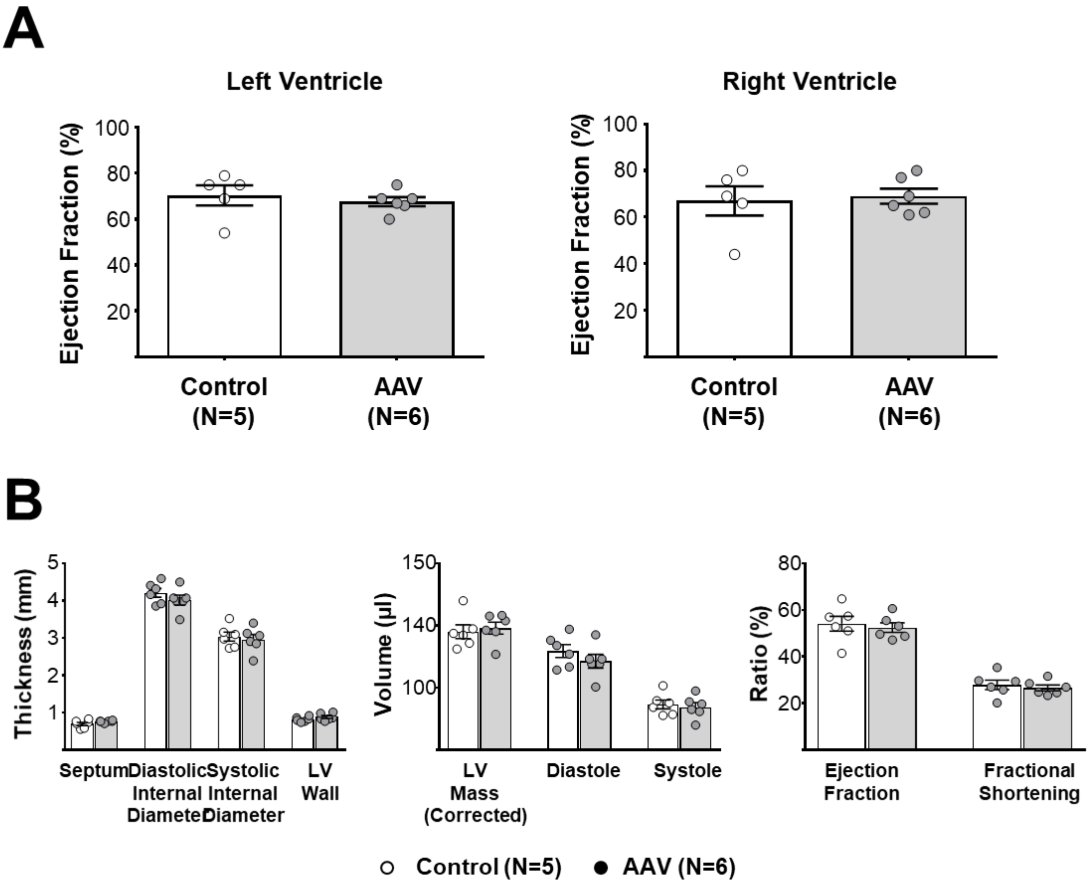


**Online Figure II. AAV - Kir2.1$^{\Delta 314-315}$ expression does not modify cardiac function analyzed by MRI and echocardiography. A.** Quantification of ejection fraction (EF), stroke volume (SV), endsystolic volume (ESV) and enddiastolic volume (EDV) in left (top) and right ventricle (bottom) by magnetic resonance imaging (MRI) in control (WT) and AAV-transduced anesthetized mice. **B.** Quantification of ejection fraction (EF), stroke volume (SV), endsystolic volume (ESV) and enddiastolic volume (EDV) in left (top) and right ventricle (bottom) by cardiac echocardiography in control (WT) and AAV-transduced anesthetized mice. * = p<0.05.

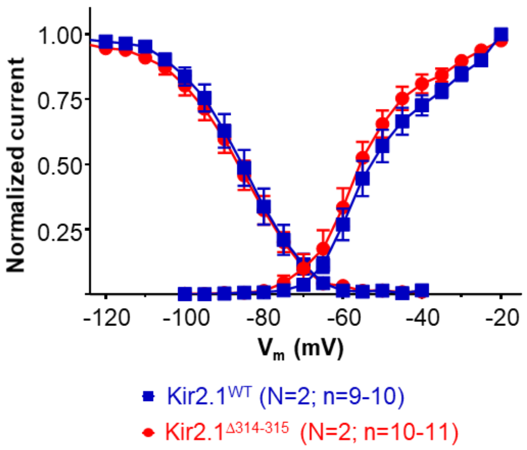


***Activation***

| | Kir2.1$^{WT}$ | Kir2.1$^{\Delta 314\text{-}315}$ |
|---|---|---|
| $V_{1/2}$ | -51.9 ± 2.45 | -53.29 ± 1.25 |
| s | 7.52 ± 1.13 | 5.34 ± 0.46 |

***Inactivation***

| | Kir2.1$^{WT}$ | Kir2.1$^{\Delta 314\text{-}315}$ |
|---|---|---|
| $V_{1/2}$ | -84.34 ± 1.85 | -83.84 ± 1.31 |
| s | 6.43 ± 0.18 | 6.76 ± 0.17 |

**Online Figure III. Cardiac expression of AAV-Kir2.1$^{\Delta 314\text{-}315}$ does not modify *$I_{Na}$* voltage-dependence of activation or inactivation.** *Left,* Na channel steady-state activation and inactivation curves for AAV-Kir2.1$^{WT}$ and AAV-Kir2.1$^{\Delta 314\text{-}315}$ cardiomyocytes plotted according to the protocol detailed in the Methods section. Currents were plotted as a fraction of peak current and fitted according to a Boltzmann function. *Right,* table shows the activation and inactivation parameters for both experimental groups.

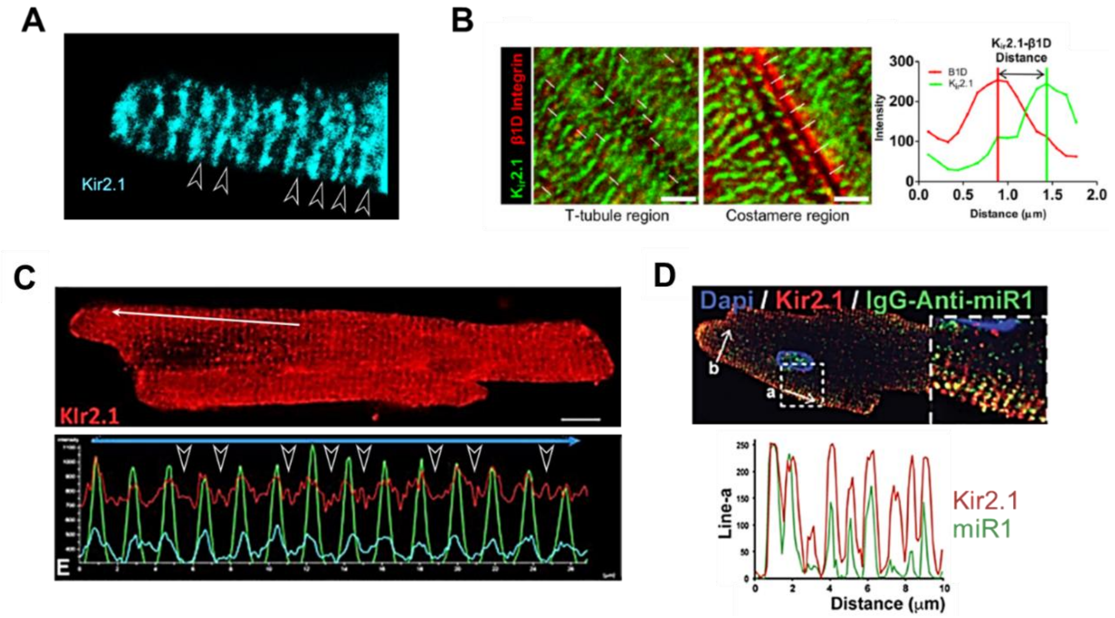


**Online Figure IV. Double Kir2.1 band staining pattern in isolated cardiomyocytes in the literature.** Presence of Kir2.1 at the SR is suggested in previously published immunostaining images, as follows: **A**, Inter-tubular Kir2.1 staining (white arrowheads) in isolated rat cardiomyocytes (from Ponce-Balbuena et al. 2018[2]); **B**, Magnified images of t-tubule (*left*) and costamere (middle) regions stained for Kir2.1 with ß1D integrin as t-tubule marker. Representative green and red fluorescence intensity profiles along one of the white lines shown on left and middle. Note the presence of Kir2.1 bands intercalated with those of ß1D (from Sengupta et al. in 2019)[12]; **C**, Immunostaining (*top*) and traces of fluorescence intensity spatial profiles (*bottom*) of Kir2.1 and miR1 through 'a' arrow-line in immunostaining. **Note the presence of one line every ≈1µm** (from Yang et al. in 2021)[13]. **D**, Inter-tubular Kir2.1 staining (white arrowheads) in isolated rat cardiomyocytes (from Park et al. in 2020[14]). All figures reproduced by permission of the respective authors and journals.

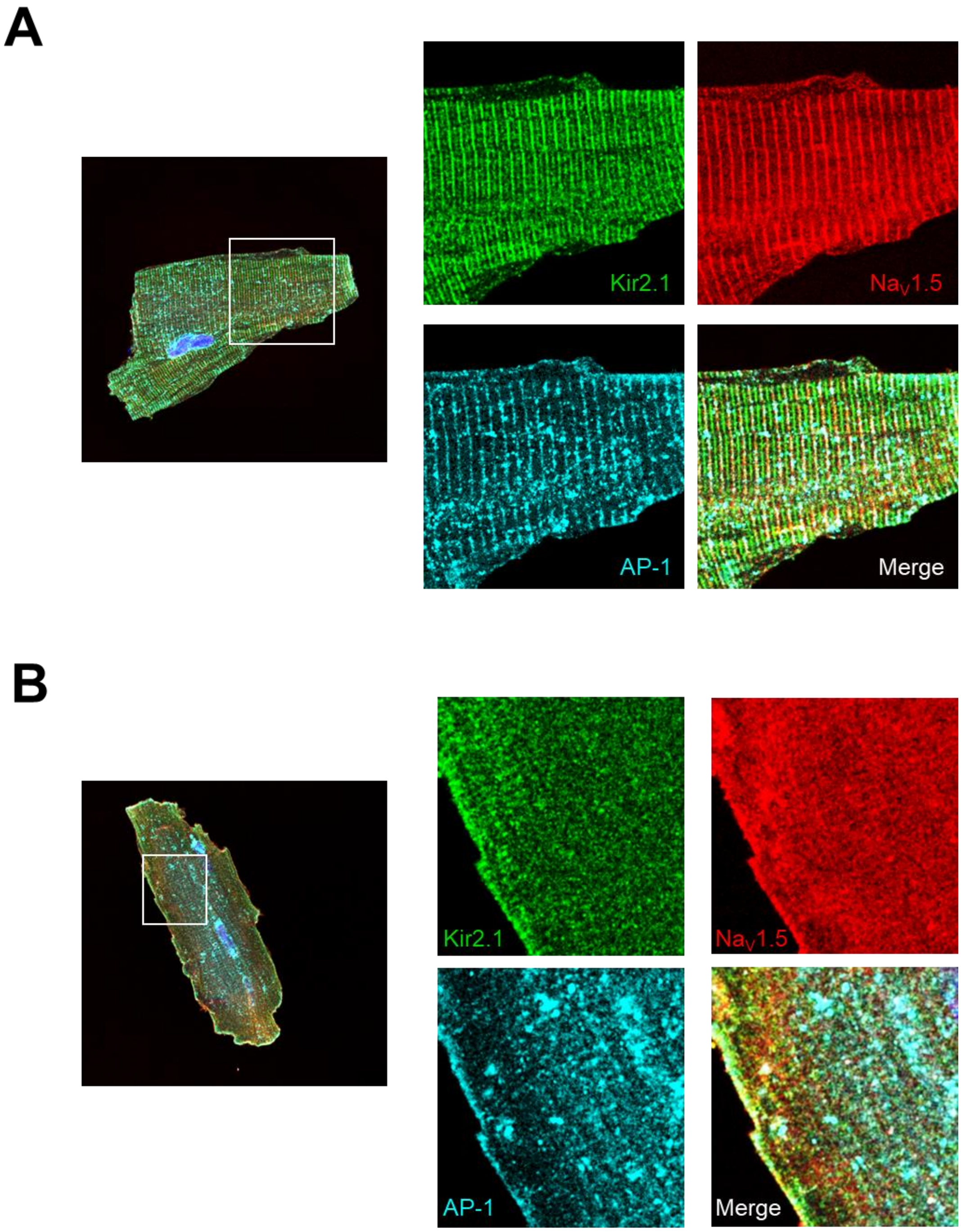


**Online Figure V. Mislocalization of Adaptin in Kir2.1$^{\Delta314\text{-}315}$ cardiomyocytes likely contributes to Nav1.5 mislocalization.** Confocal images show the expression pattern of Adaptin (AP1), Nav1.5 and Kir2.1 in Kir2.1$^{WT}$ (**A**) and Kir2.1$^{\Delta314\text{-}315}$ (**B**) cardiomyocytes.

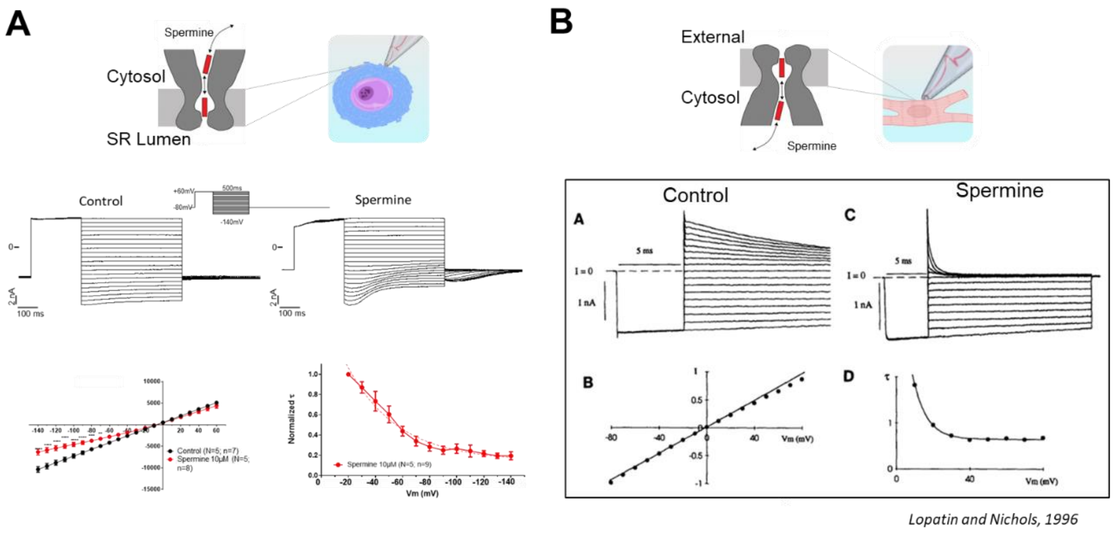


**Online Figure VI. Inverted polarity of spermine effect is likely due to the flipped channel orientation in the membrane. A. Top.** Schematic diagram of expected orientation of SR Kir channel with respect to SR lumen and cytosol in a nuclear membrane experiment. **Middle.** Representative recordings of SR Kir2.1 currents in control and in the presence of spermine. **Bottom,** Current voltage relationship and normalized SR Kir2.1 current (same data as in **Figure 6C). B. Top.** Schematic diagram of expected orientation of sarcolemmal Kir channel as reported by Nichols and Lopatin (1996)[15]. **Middle.** Representative recordings of sarcolemmal Kir2.1 currents in control and in the presence of spermine. **Bottom,** Current voltage relationship and normalized sarcolemmal Kir2.1 current. Note that differences between data in **A** and **B** are likely related to differences in channel orientation within each lipid bilayer.
